# multiMiAT: An optimal microbiome-based association test for multicategory phenotypes

**DOI:** 10.1101/2022.06.28.497893

**Authors:** Han Sun, Yue Wang, Zhen Xiao, Xiaoyun Huang, Haodong Wang, Tingting He, Xingpeng Jiang

## Abstract

Microbes affect the metabolism, immunity, digestion and other aspects of the human body incessantly, and dysbiosis of the microbiome drives not only the occurrence but also the development of disease (i.e., multiple statuses of disease). Recently, microbiome-based association tests have been widely developed to detect the association between the microbiome and host phenotype. However, existing methods have not achieved satisfactory performance in testing the association between the microbiome and ordinal/nominal multicategory phenotypes (e.g., disease severity and tumor subtype). In this paper, we propose an optimal microbiome-based association test for multicategory phenotypes, namely, multiMiAT. Specifically, under the multinomial logit model framework, we first introduce a microbiome regression-based kernel association test (multiMiRKAT). As a data-driven optimal test, multiMiAT then integrates multiMiRKAT, score test and MiRKAT-MC to maintain excellent performance in diverse association patterns. Massive simulation experiments prove the excellent performance of our method. multiMiAT is also applied to real microbiome data experiments to detect the association between the gut microbiome and clinical statuses of colorectal cancer development and the association between the gut microbiome and diverse development statuses of *Clostridium difficile* infections.

## Introduction

As a hot topic in the past decade, human microbiome studies have revealed the influence of the microbiome on physiology, metabolism, disease, and cancer [1–5]. The human microbiome inhabits various organs of the human body, including the respiratory tract, mouth, stomach, gut, skin and genitourinary tract [6]. The number of microbes is estimated to be more than 10 times the number of human cells [7]. Microbes in different parts of the human body show significant differences in microbiome composition, leading to unique microbial signatures [8]. For example, gut microbiota can produce vitamins and secrete neurotransmitters to benefit human health [9,10]. Conversely, the dysbiosis of microbial communities may be related to diseases or cancers, such as type 2 diabetes [11], obesity [12] and colorectal cancers [4]. In addition, microbes not only influence the occurrence of disease but also may have a significant impact on the development of disease. Recently, more studies have involved the role of microbiome dysbiosis in disease development, for example, disease severity [13, 14] and tumor subtype [15]. The association between disease development and the microbiome is a fundamental requirement for understanding disease mechanisms and microbes to better carry out microbial adjuvant treatment for diseases, such as targeting microbial foods [16, 17].

High-throughput sequencing (HTS) technology has made great progress in the identification and characterization of the microbiome due to its advantage of low cost [18], where amplicon and metagenomic sequencing are common HTS methods for microbiome studies [19]. Standard methods, such as QIIME [20], are usually utilized to cluster sequencing data into operational taxonomic units (OTUs) according to sequence similarity [21], where each OTU as a microbiota feature can be mapped to a species. A wide variety of OTUs are aggregated at the kingdom, phylum, class, order, family, genus, and species levels according to evolutionary relationships [22]. Overall, the features of microbiome data are a high-dimensional and phylogenetic relationship. Microbial studies usually perform global hypothesis tests to explore the association between the microbiome and host phenotype, where the type of phenotype is determined by the distribution it follows, for example, Gaussian (BMI), Bernoulli (health/disease) and multinomial distribution (health/moderate disease/severe disease). Adonis [23] and ANOSIM [24, 25] are classic distance-based statistical methods widely used to detect the association between the microbiome and host phenotypes in most microbial studies [26–29]. However, the disadvantage of these methods is that they do not adequately consider the effects of confounding factors, resulting in power loss. In addition, these distance-based methods can only adopt a single distance, and the inappropriate distance setting is also underpowered. Therefore, the influence of confounding factors and setting the distance are major challenges for these classic statistical methods [30].

Recently, microbiome-based association tests have been widely developed [31–34] to detect the association between the microbiome and host phenotypes. These methods conduct a regression model to incorporate confounding factors and the microbiome in connection with host phenotypes. For example, the optimal microbiome regression-based kernel association test (OMiRKAT) [31] incorporates the similarity metrics between samples into the regression model. A series of OMiRKAT-based methods have been extended for diverse types of outcomes [35], such as survival data [36, 37] and longitudinal data [38, 39]. These methods acquire great statistical power in testing the association between the microbiome and outcome. In addition, a number of statistical methods for microbiome-based association tests have been developed for different association patterns (i.e., sparsity level and phylogenetic relevance of association signals) [32–34, 40], such as sparse association signals [34, 40]. However, most existing microbiome-based association tests are conducted to deal with outcomes obeying Gaussian, Bernoulli, or Poisson distribution (Table 1).

**Table 1.** The scopes of use for microbiome-based association tests. These methods are presented in chronological order of publication. # represents multivariate outcomes and * denotes longitudinal data.

|  | Method | Outcome type |  |  |  |  | Reference |
| --- | --- | --- | --- | --- | --- | --- | --- |
|  |  | Continuous | Binary | Count | Survival | Multicategory |  |
| Previous Methods | OMiRKAT | ✓ | ✓ |  |  |  | [31] |
|  | aMiSPU | ✓ | ✓ |  |  |  | [32] |
|  | MMiRKAT | ✓ # |  |  |  |  | [41] |
|  | MiRKAT-S |  |  |  | ✓ |  | [36] |
|  | KRV | ✓ # |  |  |  |  | [42] |
|  | OMiAT | ✓ | ✓ |  |  |  | [43] |
|  | OMiSA |  |  |  | ✓ |  | [37] |
|  | CSKAT | ✓ * |  |  |  |  | [38] |
|  | aMiAD | ✓ | ✓ |  |  |  | [33] |
|  | aGLMMMiRKAT | ✓ * | ✓ * | ✓ * |  |  | [39] |
|  | MiHC | ✓ | ✓ | ✓ |  |  | [34] |
|  | MiATDS | ✓ | ✓ |  |  |  | [40] |
|  | MiRKAT-IQ | ✓ |  |  |  |  | [44] |
|  | AMAT | ✓ | ✓ |  |  |  | [45] |
|  | RFtest | ✓ | ✓ |  |  |  | [46] |
|  | MiRKAT-MC |  | ✓ |  |  | ✓ | [47] |
|  | aGEEMiHC | ✓ * | ✓ * | ✓ * |  |  | [48] |
| Our Method | multiMiAT |  | ✓ |  |  | ✓ |  |

The multicategory phenotypes are classified into ordinal and nominal multicategory phenotypes according to the relationship among the host phenotypes [49]. There is a progressive relationship between adjacent pairs of ordinal multicategory phenotypes, such as disease severity [13, 14], but not in nominal multicategory phenotypes, for example, tumor subtype [15] or dietary pattern [50]. Researchers often turn to binary problems to meet the requirements of existing methods when dealing with multicategory phenotypes, and two common strategies of transformation are widely used. One strategy is to choose a dividing line for classification [51]. Specifically, two contiguous statuses of ordinal outcomes are usually divided into one category, for example, genome-wide association study of COVID-19 severity [52]. The other strategy is all pairwise comparisons, for example, the study of coronary artery disease severity [53] or gut microbiome transition of four Himalayan populations [54]. Recently, a proportional odds logistic mixed model (POLMM) [51] and subtype analysis with somatic mutations (SASOM) [55] have been conducted for multicategory phenotypes. However, these methods may lose power when analyzing microbiome data because they do not consider the characteristics of microbiome data (i.e., high-dimensional and phylogenetic relationships). The microbiome kernel association test with multi-categorical outcomes (MiRKAT-MC) [47], another extension of OMiRKAT, has good performance for multicategory phenotypes, but this method is not very satisfying in some scenarios, especially nominal multicategory phenotypes.

We propose an optimal microbiome-based association test for multicategory phenotypes, namely, multiMiAT. Considering the excellent performance for OMiRKAT and its extensions, we first introduce the microbiome regression-based kernel association test (multiMiRKAT). We take inspiration from the association test for multivariant phenotypes [56] to establish a kernel machine multinomial regression framework via multinomial logit models [57]. Specifically, we conducted the connection between random component (i.e., ordinal/nominal multicategory phenotypes) and systematic component (i.e., microbiome profiles and confounding factors) based on two multinomial logit models (i.e., cumulative link model and baseline category logit model). To fully extract complex associations between the microbiome and host phenotypes, we construct numerous similarity matrices between samples through diverse distances and kernel functions. Then, we use the model fitting and similarity matrices to model the test statistic of individual tests. multiMiRKAT integrates these individual tests. Unlike MiRKAT-MC, our method focuses more on differences in host phenotypes. Because of the difference in modeling, multiMiRKAT is more advantageous in nominal multicategory phenotypes. Compared with all extensions of OMiRKAT, multiMiRKAT contains more nonlinear kernel functions (i.e., Gaussian kernel and Laplacian kernel [58]). To further capture the association signals with diverse association patterns, multiMiAT integrates the bcl-based score test [55] and MiRKAT-MC on the basis of multiMiRKAT. Extensive simulation experiments demonstrate that our method has excellent statistical power with the correct type I error rates. We apply multiMiAT to two real microbiome datasets. An efficient and user-friendly R package **multiMiAT** is available at https://github.com/xpjiang-ccnu/multiMiAT.

## Materials and methods

In this section, we describe our methods in detail, including multinomial logit models, test statistics and *p* value calculation (Figure 1). Specifically, we first adopt two multinomial logit models to connect the association between the marginal mean of multicategory phenotypes and the systematic component, where the systematic component contains microbiome profiles and confounding factors. We construct many similarity matrices between subjects using seven distances and three kernel functions. Then, we perform the test statistic of the microbiome regression-based kernel individual test via the model fitting and similarity matrices. We integrate these individual tests to establish multiMiRKAT (i.e., multiMiRKAT-O and multiMiRKAT-N). Finally, we model multiMiAT by integrating the multiMiRKAT, score test and MiRKAT-MC to ensure strong power in diverse scenarios.

**Figure 1.**
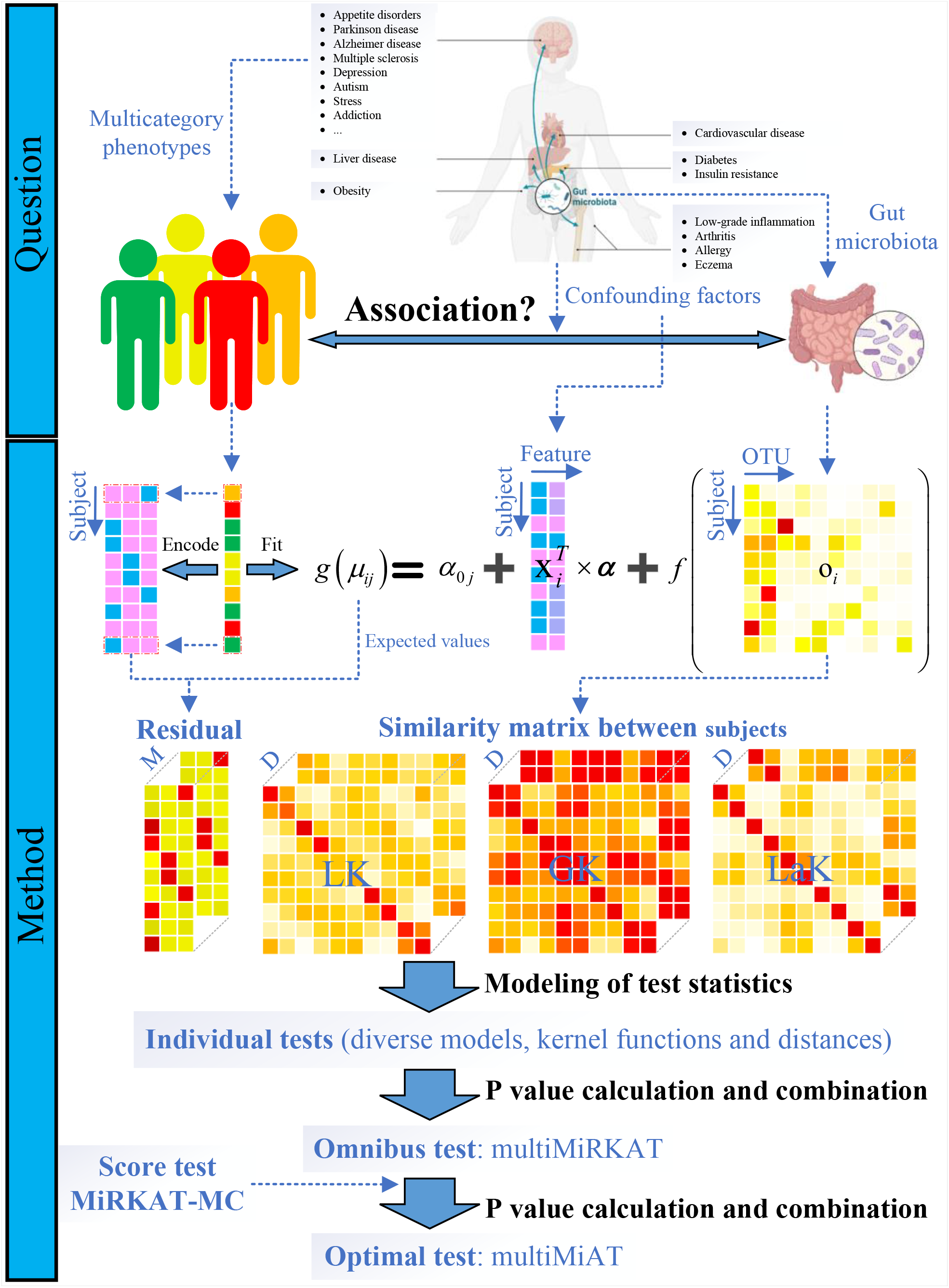
Schematic depiction of the framework for multiMiAT. Our method can be used for detecting the association between microbiome and multicategory phenotypes. Specifically, we first establish microbiome regression-based kernel individual tests via multinomial logit models. Then, we conduct the omnibus test (multiMiRKAT) through integrating these individual tests. To accommodate diverse association patterns, multiMiAT integrates multiMiRKAT, bcl-based score test and MiRKAT-MC.

### Multinomial logit models

For the observation value of the *i*th subject, we record the multicategory outcomes *y*_*i*_ ∈ {1, 2, …, *J* } (i.e., the multicategory phenotypes of interest), where *i* = 1, 2, …, *n* and *J* ≥ 2. We suppose *Y*_*ij*_ = I (*y*_*i*_ = *j*), where I (·) is the indicator function and *j* = 1, …, *J* − 1, and define **Y**_*i*_ = (*Y*_*i*1_, …, *Y*_*i*(*J*−1)_)^*T*^. We also observe *s* covariates **x**_*i*_ = (*x*_*i*1_, …, *x*_*is*_)^*T*^ and the abundance **o**_*i*_ = (*o*_*i*1_, *o*_*i*2_, …, *o*_*im*_)^*T*^ of *m* OTUs. We assume that the marginal expectation of *Y*_*ij*_ given **x**_*i*_, **o**_*i*_ is *µ*_*ij*_ := *E*(*Y*_*ij*_|**x**_*i*_, **o**_*i*_) = *P* (*y*_*i*_ = *j*|**x**_*i*_, **o**_*i*_). We connect the marginal mean of multicategory phenotypes and systematic component, including covariates and microbiome profiles, by specifying that

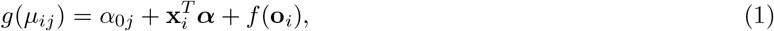

where *α*_0*j*_ and ***α*** are the intercept and regression coefficients, respectively. *g*(·) is a link function. *f* (·) is a function reflecting the microbiome profile. The popular approach is a positive definite kernel function *K*(·,·), that is 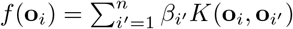 [31, 36].

Multicategory outcomes are usually divided into ordinal or nominal multicategory outcomes according to the relationship between categories [57]. The progressive relationship occurs between any adjacent statuses of ordinal multicategory outcomes, such as the degree of disease severity (health/mild/moderate/severe), and this effect is cumulative and always increased to the maximum at both extreme statuses. For ordinal multicategory outcomes, we adopt the cumulative link model (clm, also called the proportional odds model) as the link function, whose form is as follows:

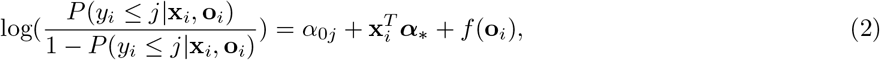

where the regression coefficients ***α***_∗_ are fixed for all *j* in Eq. 2. *α*_0*j*_ is the category-specific intercept and needs to satisfy a monotonicity constraint (i.e., *α*_01_ ≤ … ≤ *α*_0(*J*−1)_) only in the cumulative link model [59].

For the nominal multicategory outcomes, there is no progressive relationship between the statuses of nominal outcome. We utilize the baseline category logit model (bcl) as the link function, whose form is as follows:

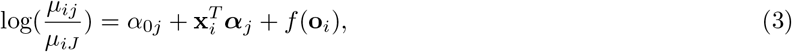

where ***α***_*j*_ is category-specific regression coefficient in Eq. 3. Our goal is to test the association between the microbiome profiles and host phenotypes. We define the variance of microbiome profiles *f* (**o**_*i*_) is *τ*. We assume that there is no association between OTUs and host phenotypes as the global null hypothesis, that is, *H*_0_: *τ* = 0.

### Similarity matrix between subjects

The dissimilarity matrix between subjects is used to measure a between-group difference in microbiome data, such as Adonis and ANOSIM [24,25]. The similarity matrix is usually obtained by transforming the dissimilarity matrix between subjects through the kernel function. Due to weight assignments (unweighted or weighted) and covered information (with or without phylogeny information), the forms of the dissimilarity vary. Different dissimilarities take disparate weight assignments, for example, more weight assigned either to rare lineages or to most abundant lineages, may lead to inconsistent results in the dissimilarity-based analysis. To ensure fairness in comparative analysis, we adopt the same distance configuration as OMiRKAT. Specifically, Bray-Curtis dissimilarity [60] (D_BC_), a non-phylogeny-based dissimilarity, and unweighted and weighted UniFrac [61, 62] (D_u_ and D_w_), the phylogeny-based distances, are taken into account because they are also commonly used in microbial research. Considering weight assignments, we also adopt the four generalized UniFrac [63] with *α* = 0, 0.25, 0.5, 0.75 (D_0_, D_0.25_, D_0.5_, D_0.75_).

To measure the similarity between subjects from their microbiome profiles, we convert the distance matrix into a kernel matrix via a kernel function. Most previous kernel-based regression microbiome association analyses only consider the linear kernel (K_LK_) function [31, 39, 41, 43], whose form is as follows:

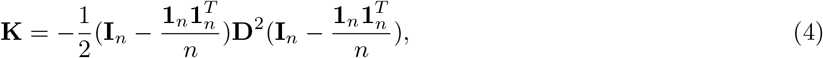

where **I**_*n*_ is a *n* × *n* identity matrix and **1**_*n*_ is a *n*-dimensional vector of ones. **D** is the dissimilarity matrix. However, the linear kernel may struggle with more complex associations. Thus, we consider taking more nonlinear kernel functions into account in our model, such as the Gaussian kernel function and Laplacian kernel function [58]. The models are as follows:

1. Gaussian kernel (K_GK_) function: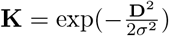,
2. Laplacian kernel (K_LaK_) function: :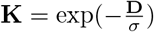,

where *σ* is the hyperparameter. Here, we adopt *σ* = 1 after a large amount of experimental verification (Supplementary Figure S1). We also conduct the positive semidefiniteness correction procedure for **K** to ensure that the eigenvalue of **K** is nonnegative [31]. We calculate the eigenvalues *λ*_*i*_ (*i* = 1, …, *n*) and eigenvectors of **K** via eigenvalue decomposition **K** = **FΛF**^*T*^, where **Λ** is the diagonal matrix with the *i*th diagonal element *λ*_*i*_. Then, we obtain the reconstructed kernel matrix **K**^∗^ = **FΛ**^∗^**F**^*T*^, where **Λ**^∗^ is the diagonal matrix with the *i*th diagonal element |*λ*_*i*_|. Here, we assume that 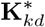 represents a similarity matrix obtained by using diverse kernel functions and dissimilarity measures, where *k* ∈ Γ_K_ = {K_LK_, K_GK_, K_LaK_} and *d* ∈ Γ_D_ = {D_BC_, D_u_, D_0_, D_0.25_, D_0.5_, D_0.75_, D_w_}.

### Microbiome regression-based kernel individual tests

To detect the association signals between microbiome profiles and host phenotypes, we need to establish our test statistics and calculate the *p* values. Compared with the test statistics for the binary phenotype, the residual in test statistics for the multicategory phenotypes is not *n*-dimensional but *n* × (*J* − 1)-dimensional. Thus, it is a challenge to extend the original model to our model. Inspired by the association test for multivariant phenotypes [56], we establish our test statistics of individual tests

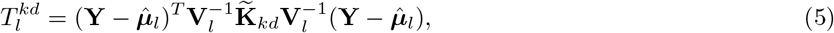

where 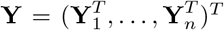 and 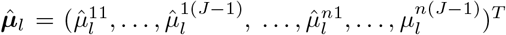. 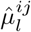 is the estimated expectation via multinomial logit model *l* under the *H*_0_ condition, where *l* ∈ Γ_L_ = {M_bcl_, M_clm_}. **V**_*l*_ = **I**_*n*_ ⊗ **V**_*l*0_ and **V**_*l*0_ is the estimated residual variance matrix under the null model. Here, ⊗ is the Kronecker product. 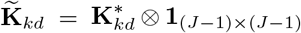, where **1**_(*J*−1)×(*J*−1)_ is a (*J* − 1) × (*J* − 1) matrix of ones. Obviously, the original test statistic (i.e., OMiRKAT) for the binary phenotype is a special case (i.e., dealing with binary outcome) of our test statistics 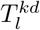.

### Microbiome regression-based kernel omnibus tests

The setting of distance and kernel function can affect similarity between samples and further lead to performance differences among individual tests because of the weight assignments or covered information for different distances. For the situation where the association signals have a phylogenetic relationship, the distance covering phylogenetic tree information should be used. When the differences among different phenotypes are mainly in the presence/absence or abundance of microbial communities with phylogenetic relationships, the unweighted or weighted distances may be more appropriate, respectively. In addition, the association between microbiome profiles and multicategory phenotypes may not be linear. Considering the complexity of association patterns, we take the minimum *p* values of individual tests, namely, MinP, as the test statistics of global omnibus tests

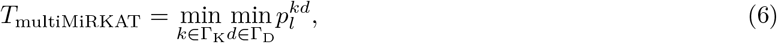

where 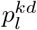 denotes the *p* value of 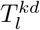. MinP is widely used in microbiome-based association tests, such as OMiRKAT [31], OMiAT [43] and MiHC [34]. For ease of comparison, we define the test statistic of the local omnibus test 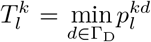, which integrates individual tests with diverse distances. Obviously, the local omnibus test with a linear kernel function is OMiRKAT based on a multinomial logit model. In addition, considering that the regression coefficients fitted by the ordinal/nominal multinomial logit model may be inconsistent, we adopt two multinomial logit models (i.e., bcl and clm) to model the global omnibus tests. We call the bcl-based and clm-based global omnibus test multiMiRKAT-N and multiMiRKAT-O, respectively.

multiMiRKAT considers a variety of individual tests with diverse distances, kernels and models, which may put considerable pressure on the calculation of the *p* value. Thus, we fully consider the configuration of the distance and kernel function to ensure computational efficiency. For example, we discuss the performances of other distance (i.e., Jaccard dissimilarity [64]), kernel function (i.e., exponential kernel function) and ordinal multinomial logit model (i.e., continuation ratio model, crm). In addition, we also explore other *p* value combination methods, such as the aggregated Cauchy association test (ACAT) [65] and the harmonic mean *p*-value (HMP) [66]. For specific comparison results, please refer to the section “Components affecting performance”.

### Score test

Schaid et al. [67] introduced the GLM-based score test, which is used for detecting the association between traits and haplotypes. Then, the extended versions of the score test were widely developed, for example, the GEE-based score test [68] and bcl-based score test [55,69]. Here, we consider integrating the bcl-based score test into our method to improve the statistical power for detecting abundant association signals. We define 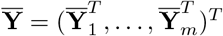 and **O** = (**o**_1_, …, **o**_*n*_)^*T*^, where 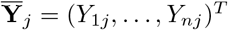. **W** represents a *m*-dimensional weight vector that represents the weights of *m* OTUs. Here, we adopt **W** = **1**_*m*_ and **1**_*m*_ is a vector of ones. Let **S** = **I**_(*J*−1)_ ⊗ (**OW**^*T*^), where **I**_(*J*−1)_ represents a (*J* − 1) × (*J* − 1) identity matrix. The covariance matrix of 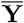 is

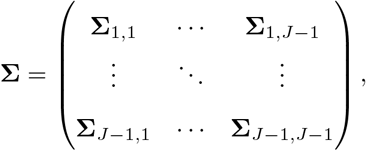

where

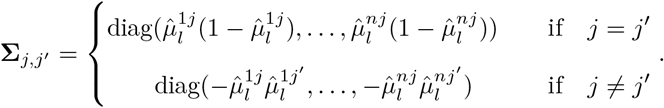

We define 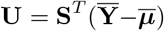 and X = **I**_(*J*−1)_⊗**X**, where **X** = (**x**_1_, …, **x**_*n*_)^*T*^ and 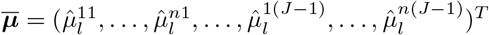. Then, we establish the test statistic of the score test

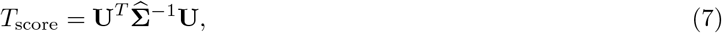

where 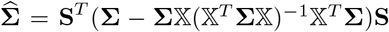 is the covariance matrix of **U**. *T*_score_ can be shown to asymptotically follow a *χ*^2^ distribution with degree of freedom (*J* − 1) [55].

### MiRKAT-MC

MiRKAT-MC [47] is proposed for multicategory phenotypes. Similar to OMiRKAT and all its extensions, MiRKAT-MC also utilizes the *p* value combination method (i.e., HMP) to integrate individual tests. Here, the test statistic of the individual test is

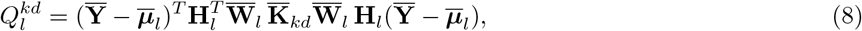

where 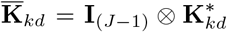 and 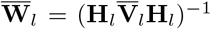. 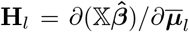 and 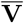 is the variance-covariance matrix. The *p* values of individual tests are calculated via a pseudo-permutation strategy [42]. MiRKAT-MC uses the HMP to combine these *p* values of individual tests, where MiRKAT-MCN and MiRKAT-MCO are bcl-based and clm-based MiRKAT-MC, respectively. Obviously, multiMiRKAT is different from MiRKAT-MC in the modeling of test statistics. MiRKAT-MC focuses more on differences in the similarity matrices rather than host phenotypes, resulting in good performance in the rare association signals. Remarkably, compared with the original MiRKAT-MC, we consider more distances and kernel functions in our models to enhance the performance of MiRKAT-MC in diverse association patterns.

### Optimal test for multicategory outcomes

In fact, the true association patterns are usually unknown and difficult to predict. Due to differences in modeling, the above three methods (i.e., multiMiRKAT, score test and MiRKAT-MC) have their own advantages in different scenarios. Specifically, multiMiRKAT and MiRKAT-MC perform well for detecting association signals with phylogenetic relationships because a large number of distances covering phylogenetic information are used. multiMiRKAT takes the idea of dealing with multivariate phenotypes, and compared with MiRKAT-MC, multiMiRKAT may perform better for nominal multicategory phenotypes. The score test is sensitive to abundant association signals because its test statistic is established based on the model fitting only. Overall, a single method does not always perform best in all scenarios. Therefore, we further present multiMiAT by integrating these three methods. Since the score test and MiRKAT-MC do not use the permutation method to calculate their *p* values, we only adopt HMP to combine the *p* values and establish the test statistic of multiMiAT

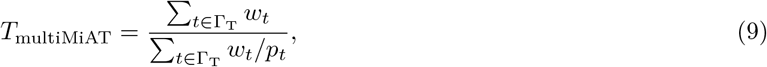

where Γ_T_= {*T*_multiMiRKAT-N_, *T*_multiMiRKAT-O_, *T*_score_, *T*_MiRKAT-MCN_, *T*_MiRKAT-MCO_} and 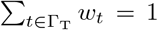. *p*_*t*_ represents the *p* values of these combined tests. Considering the differences in modeling, these methods may be complementary in some scenarios, which further enables the optimal test to maintain excellent performance in diverse association patterns.

### *P* values calculation

We consider the methods based on permutation and distribution to conduct the *p* value calculation, which are common in *p* value analysis [31, 66]. Specifically, we perform the *p* value calculation of multiMiRKAT, including individual tests and omnibus tests, via the permutation method. Because we utilize the HMP method to establish the test statistic of multiMiAT, the method based on distribution is used for the *p* value of multiMiAT. The detailed calculation process can be found in the supplementary material.

### Comparison methods

In the comparative analysis section, we consider two types of methods: association tests for multicategory outcomes and microbiome-based association tests for binary outcomes. Here, the association tests for multicategory outcomes include Adonis [23], ANOSIM [24, 25], SASOM [55] and MiRKAT-MC [47]. Specifically, Adonis and ANOSIM adopt Bray-Curtis dissimilarity. SASOM utilizes three methods to combine the *p* values of individual tests, namely, SASOM-F, SASOM-T and SASOM-D. MiRKAT-MCN and MiRKAT-MCO are bcl-based and clm-based MiRKAT-MC, respectively. These methods can be utilized for comparing the differences in microbial composition among multiple groups. The microbiome-based association tests for binary outcomes mainly include OMiRKAT [31], adaptive microbiome-based sum of powered score (aMiSPU) [32], optimal microbiome-based association test (OMiAT) [43] and microbiome higher criticism analysis (MiHC) [34], which are classic methods. Limited by being mainly designed for binary outcomes, these methods cannot directly handle multicategory outcomes. One popular strategy is taking the minimal of *J*(*J* − 1)*/*2 *p* values and conducting the product of the minimal *p* values and the number of comparisons as the results, where *J*(*J* − 1)*/*2 *p* values are obtained from all the pairwise comparisons among the *J* subtypes [55]. This strategy is Bonferroni correction for multiple hypothesis testing. For simplicity, we refer to this strategy as a pairwise analysis. However, we believe that it is not suitable for all multicategory outcomes (i.e., ordinal and nominal). We consider other strategies for ordinal and nominal multicategory outcomes, that is, adjacent and baseline pairwise analysis take the minimal of *J* − 1 *p* values and conduct the product of the minimal *p* values and *J* − 1, respectively. We perform a detailed analysis of these correction strategies in the section “Correction of microbiome-based association tests”.

## Results

### Simulation design

Our simulation experiments are designed based on previous studies [32, 36, 51, 70]. We adopt a real throat microbiome dataset [71]. For a more convenient calculation, we use the processed data, including 273 OTUs with mean relative abundance ≥ 10^−4^, which can be found in the R package *MiHC* [34]. Then, we estimate proportions and dispersion parameter of real data and use a Dirichlet-multinomial model to generate the OTU count table with 1000 total counts per sample. Let *o*_*ij*_ denote the *j*th generated OTU of the *i*th sample. Considering the effect of sample size *n*, we adopt *n* = 100 and 200. We also consider two covariates *x*_*i*1_ and *x*_*i*2_ that obey the Bernoulli distribution *B*(1, 0.5) and standard normal distribution *N*(0, 1), respectively.

We focus on testing microbial signals associated with the phenotype. As far as we know, the association pattern has a significant effect on the statistical power of association tests, where the association pattern mainly refers to the sparsity levels and phylogenetic relevance of microbiome association signals [40]. To verify the performance of our method under different association patterns, we calculate the cophenetic distances for all OTU pairs using the function, *cophenetic*, available in the R package, *stats*. We use the partitioning-around-medioids algorithm [72] to divide 273 OTUs into 15 clusters based on cophenetic distances. We design the following scenarios for the choice of microbial signals:

1. The most abundant cluster from 15 clusters consists of 49 OTUs (17.95%). The average proportion of this cluster’s reads relative to the total reads is 21.34%.
2. The mean abundant cluster from 15 clusters consists of 14 OTUs (5.13%). The average proportion of this cluster’s reads relative to the total reads is 1.43%.
3. The most rare cluster from 15 clusters consists of 5 OTUs (1.83%). The average proportion of this cluster’s reads relative to the total reads is 0.90%.
4. The 30 most abundant OTUs (10.99%) according to the total/mean reads of each OTU. The average proportion of these OTUs’ reads relative to the total reads is 60.27%.

We simulate the ordinal multicategory outcomes by using the following model:

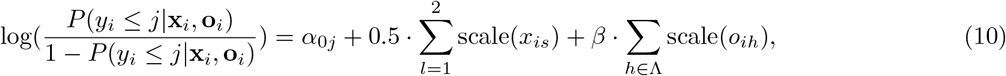

where *β* is the effect size and Λ is a set of selected OTUs. scale(·) is a zero-mean normalization function with mean 0 and standard deviation 1. Here, we only consider the number of categories *J* = 4. When generating ordinal outcomes, we also discuss the impact of balanced/unbalanced data on the results, where balanced/unbalanced data refer to the proportions of different phenotypes [51]. In particular, we cleverly perform a dynamic intercept *α*_0*j*_ to construct designed balanced and unbalanced data for ordinal phenotypes, where the dynamic intercept is obtained according to the ranking of GLM’s systematic component. Assuming the sample size of ordinal categorical phenotype including *J* = 4 levels *n*_1_, *n*_2_, *n*_3_, *n*_4_, we set *n*_1_ : *n*_2_ : *n*_3_ : *n*_4_ = 1 : 1 : 1 : 1 for balanced design and *n*_1_ : *n*_2_ : *n*_3_ : *n*_4_ = 1 : 4 : 4 : 1 for unbalanced design.

We simulate the nominal multicategory outcomes by using the following model:

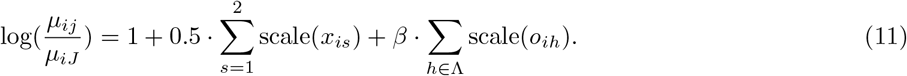

Because it is a huge challenge to obtain balanced data by setting the regression coefficients of Eq. 11, we do not design related experiments. For simplicity, we call balanced design, unbalanced design and nominal multicategory outcomes scenarios a, b, and c, respectively. To evaluate the type I error rate and power, we set *β* to control the association between host phenotypes and the microbiome. We set *β* = 0 for the type I error rate and generate 5000 simulation data repeatedly. We set *β* ≠ 0 for the power and generate 2000 simulation data repeatedly.

### Components affecting performance

In this section, we discuss the components affecting the performance of our method, that is, the hyperparameter *σ* of nonlinear kernel functions and the choice of distances, kernel functions and logit models.

For the hyperparameter *σ*, we consider *σ* ∈ {0.25, 0.5, 0.75, 1, 1.25, 1.5, M}, where “M” denotes the mean of distance matrix **D** [42]. We compare the results of their local test in scenario 1a. We find that the power is stable with *σ* = 1, but the performance of *σ* = M is poor, especially K_GK_ (Supplementary Figure S1). Therefore, we set the hyperparameter *σ* = 1.

Considering the inconsistent similarity matrices between subjects through different distances, the power of individual tests is different. For the choice of distances, we accept the distance configuration of OMiRKAT [31] because the individual tests with these distances have their own advantages in different situations (Figure 2). The performances of individual tests with phylogeny-based dissimilarity (e.g., UniFrac distance) are better for cases in which association signals with phylogenetic relationships are selected (scenarios 1-3). As the abundance of association signals decreases, the individual tests that perform better also change from individual tests based on larger weight UniFrac (D_w_, D_0.75_, D_0.5_) to individual tests based on smaller weight UniFrac (D_u_, D_0_, D_0.25_). The performance of individual tests with non-phylogeny-based dissimilarity (i.e., Bray-Curtis dissimilarity) is better for cases in which association signals without phylogenetic relationships are selected (scenario 4). As the sample size *n* increases, the performance improves (Figure 2, S2). In addition, similar conclusions can also be drawn from individual tests based on different kernel functions or logit models (Supplementary Figure S3–S5). We also consider Jaccard similarity (D_Ja_) to conduct comparative analysis with our used distances Γ_D_, where D_Ja_ is used in an extension method of OMiRKAT [39]. However, it is poor in statistical power for all scenarios (Figure 2, S2–S5).

**Figure 2.**
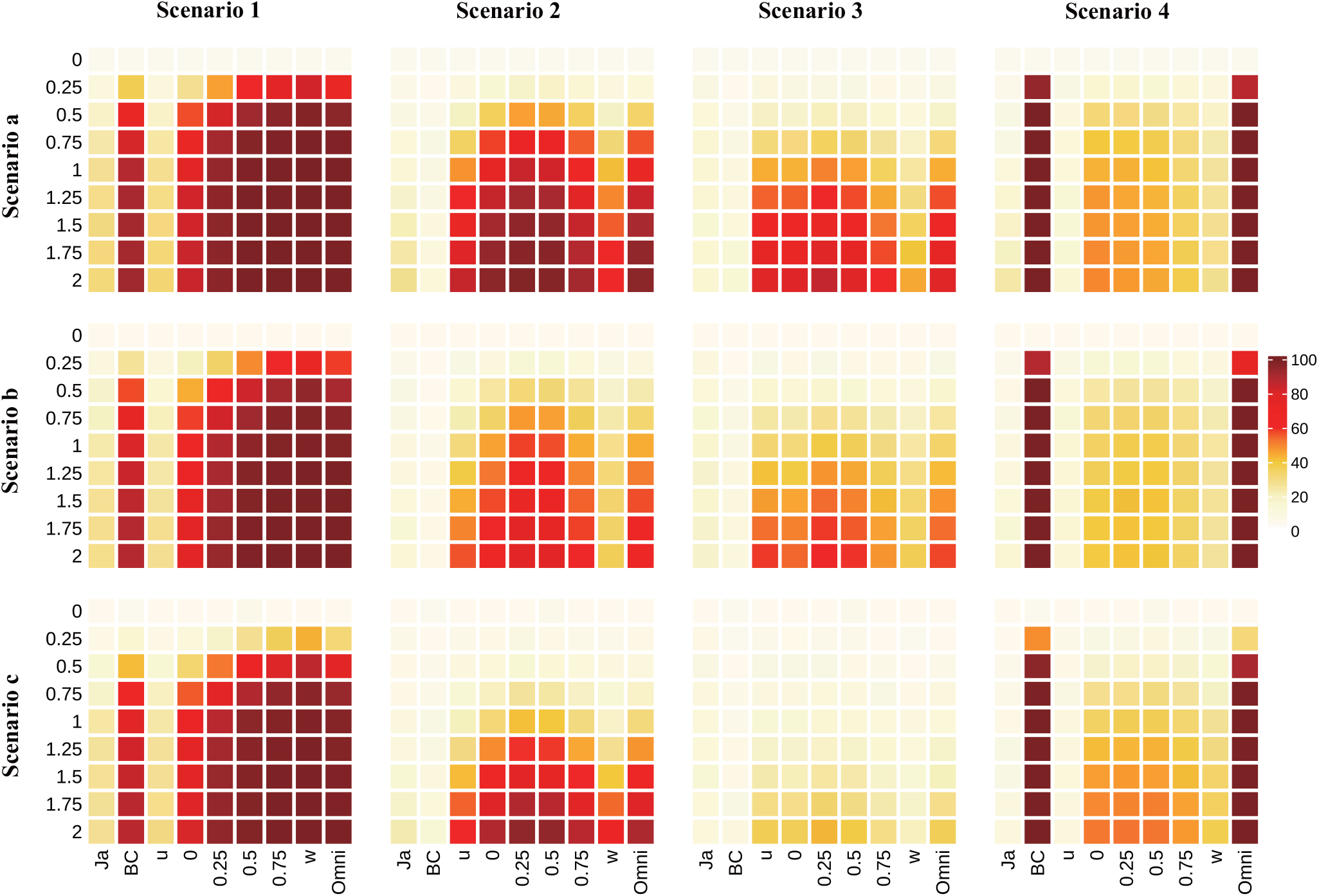
Comparison of the type I error rates and powers among microbiome regression-based kernel individual tests and local omnibus test with diverse distances under diverse scenarios (*n* = 200). Here, we adopt linear kernel and baseline category logit model.

For the choice of kernel functions, we consider the LK function, GK function, LaK function and exponential kernel (EK) function, where the form of EK is 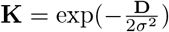. The choice of kernel function may have an impact on the power. For example, the performance of EK is slightly poorer than that of the others. In addition, for the choice of ordinal multinomial logit models, we discuss clm and crm, where the form of crm is as follows

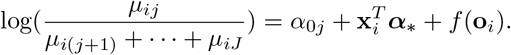

We find that the difference between clm and crm is not significant (Supplementary Figure S6). Therefore, we do not consider the EK function and crm. The local omnibus test integrating individual tests with diverse distances may not be the best in terms of statistical power, but it is always close to the best one and robust for different situations. For simulation data generated via a linear systematic component (Eq. 10, 11), nonlinear kernels can maintain performance similar to linear kernels in most cases and, in some cases, even slightly better than linear kernels (Supplementary Figure S7, S8).

In addition, we use MinP and HMP to construct the test statistics of multiMiRKAT and multiMiAT, respectively. However, the choice of *p* value combination methods may also affect the performance. We explore the performance of three different *p* value combination methods (MinP, HMP and ACAT). From the result of the empirical type I error rate and power (Supplementary Table S1–S4 and Figure S7–S10), HMP is the most conservative one, and ACAT is the most sensitive one among these methods. Specifically, the type I error rate and power of HMP are generally low. ACAT is so sensitive that it may obtain an inflated Type I error rate. Therefore, MinP may be a more suitable *p* value combination method to integrate these individual tests, and our strategy (i.e., “MinP+HMP”) has better performance compared with only using HMP.

### Correction of microbiome-based association tests

For the choice of pairwise, adjacent pairwise and baseline pairwise analysis of microbiome-based association tests, we adopt adjacent pairwise analysis for ordinal multicategory outcomes and baseline pairwise analysis for nominal multicategory outcomes in simulation experiments.

For ordinal multicategory outcomes, there is a progressive relationship between any adjacent two groups. Obviously, a cumulative effect of this progressive relationship appears among nonadjacent pairwise groups when we generate ordinal multicategory outcomes. Thus, the difference among nonadjacent pairwise groups, especially two extremely pairwise groups, is undoubtedly further amplified. In simulation experiments, we analyze the distribution of the minimum *p* values. We find that the minimum *p* values are not evenly scattered in each pairwise group but mostly in nonadjacent pairwise groups, especially in two extremely pairwise groups for the balanced design experiment (Supplementary Figure S11–S14). For unbalanced designs, the differences in nonadjacent pairwise groups are still the most significant. Although the large differences among nonadjacent groups may cause the extremely excellent performance of pairwise groups in statistical power (Supplementary Figure S15), the corrected *p* values of pairwise groups are difficult to convince anyone of the association between microbiome and all ordinal multicategory outcomes. Therefore, we consider using adjacent pairwise analysis to correct microbiome-based association tests for ordinal multicategory outcomes.

For nominal multicategory outcomes, each group is generated independently. No pairwise groups have a progressive relationship, and pairwise analysis can be utilized to correct *p* values. In the analysis of the distribution of the minimum *p* values, we find that the *p* values of pairwise groups containing the *J* th group are relatively low in most cases (Supplementary Figure S11–S14). This is probably because we use the *J* th group as the baseline group to generate the nominal multicategory outcomes. To make the strategy used more convincing, we consider baseline pairwise analysis, which always contains baseline group and one group other than baseline group and can eliminate the influence of the baseline group. In fact, the difference between baseline pairwise analysis and pairwise analysis is not significant in performance (Supplementary Figure S15). Therefore, we consider using baseline pairwise analysis to correct microbiome-based association tests for nominal multicategory outcomes.

### Type I error

The empirical type I error rates of all the methods used in the simulation experiment are reported in Table 2, S4–S6. We set the significance level *α* = 0.05, and our methods, including individual tests and omnibus tests, can be controlled approximately 5% accurately. For scenario b (i.e., unbalanced data), OMiAT obtains an inflated type I error rate because the difference among adjacent groups is more significant compared with scenario a (i.e., balanced data). As the sample size increases, this effect is alleviated.

**Table 2.** Empirical type I error rates of all methods. * represents inflated type I error rates.

| | | $n = 100$ | | | $n = 200$ | | |
| --- | --- | --- | --- | --- | --- | --- | --- |
|  | Method | Ordinal |  | Nominal | Ordinal |  | Nominal |
|  |  | Balance | Unbalance |  | Balance | Unbalance |  |
| Previous Methods | Adonis | 4.92% | 5.24% | 4.48% | 5.24% | 4.66% | 4.70% |
|  | ANOSIM | 4.82% | 4.78% | 5.08% | 5.22% | 4.94% | 4.34% |
|  | aMiSPU | 5.22% | 5.16% | 4.38% | 4.74% | 5.58% | 4.80% |
|  | MiHC | 2.92% | 5.90% | 2.56% | 3.52% | 5.32% | 3.96% |
|  | MiRKAT-MCN | 4.38% | 5.02% | 4.10% | 4.94% | 4.92% | 4.62% |
|  | MiRKAT-MCO | 4.62% | 5.44% | 4.44% | 4.24% | 5.02% | 4.50% |
|  | OMiAT | 5.02% | 7.08%* | 2.66% | 5.18% | 5.76% | 3.90% |
|  | OMiRKAT | 5.32% | 5.36% | 3.88% | 4.24% | 5.10% | 4.00% |
|  | SASOM-F | 5.40% | 5.50% | 4.94% | 5.12% | 4.62% | 4.66% |
|  | SASOM-T | 4.60% | 4.50% | 4.46% | 4.42% | 4.30% | 4.36% |
|  | SASOM-D | 5.12% | 4.98% | 4.72% | 5.06% | 4.54% | 4.52% |
| Our Method | multiMiAT | 4.68% | 5.34% | 4.72% | 4.80% | 4.96% | 4.88% |

### Power

The performances of statistical power for all the methods used in the simulation experiment are reported in Figure 3, 4, 5 and Supplementary Figure S9, S10, S16–18.

**Figure 3.**
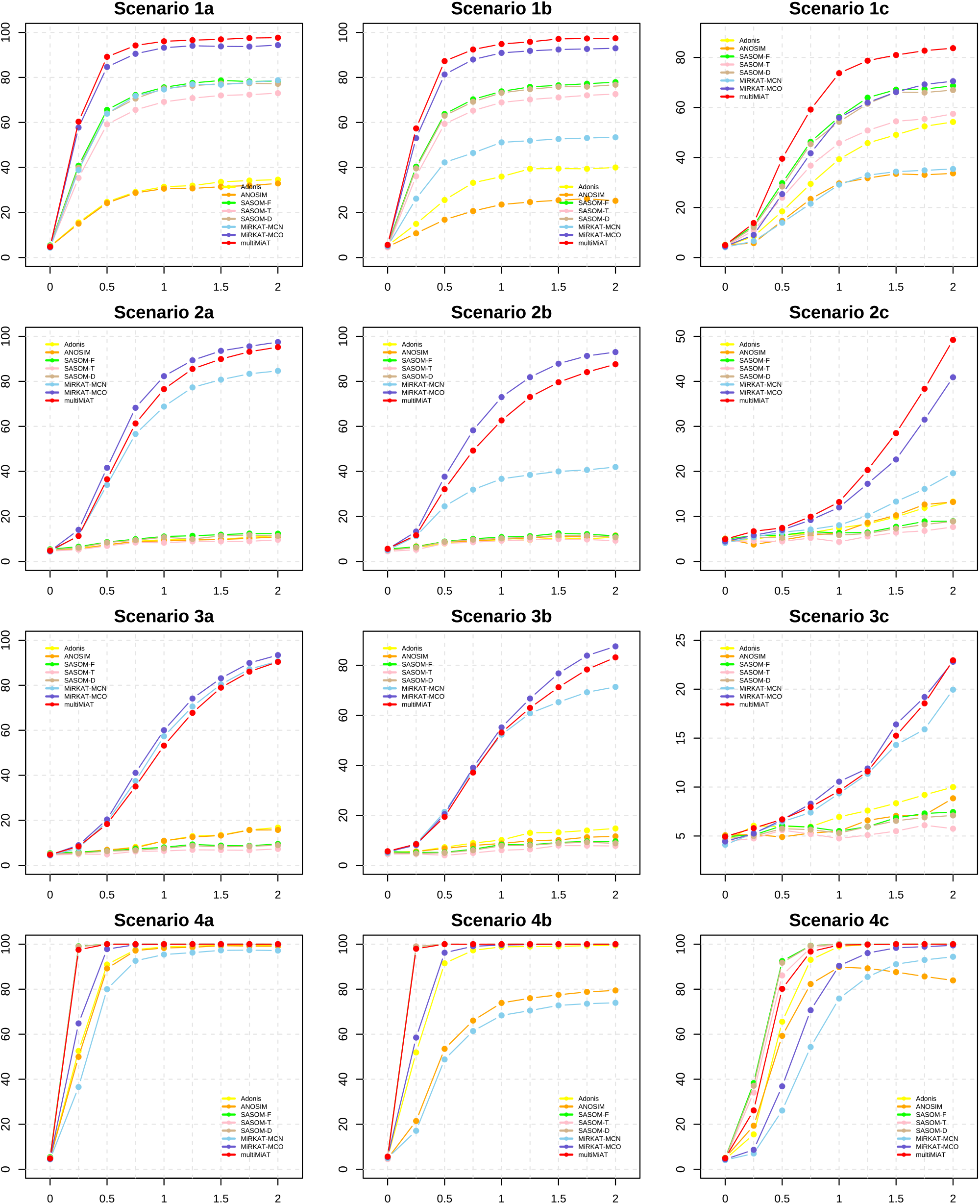
Under diverse scenarios, the comparison of the powers between our method (i.e., multiMiAT) and previous association tests for multicategory outcomes (*n* = 100).

**Figure 4.**
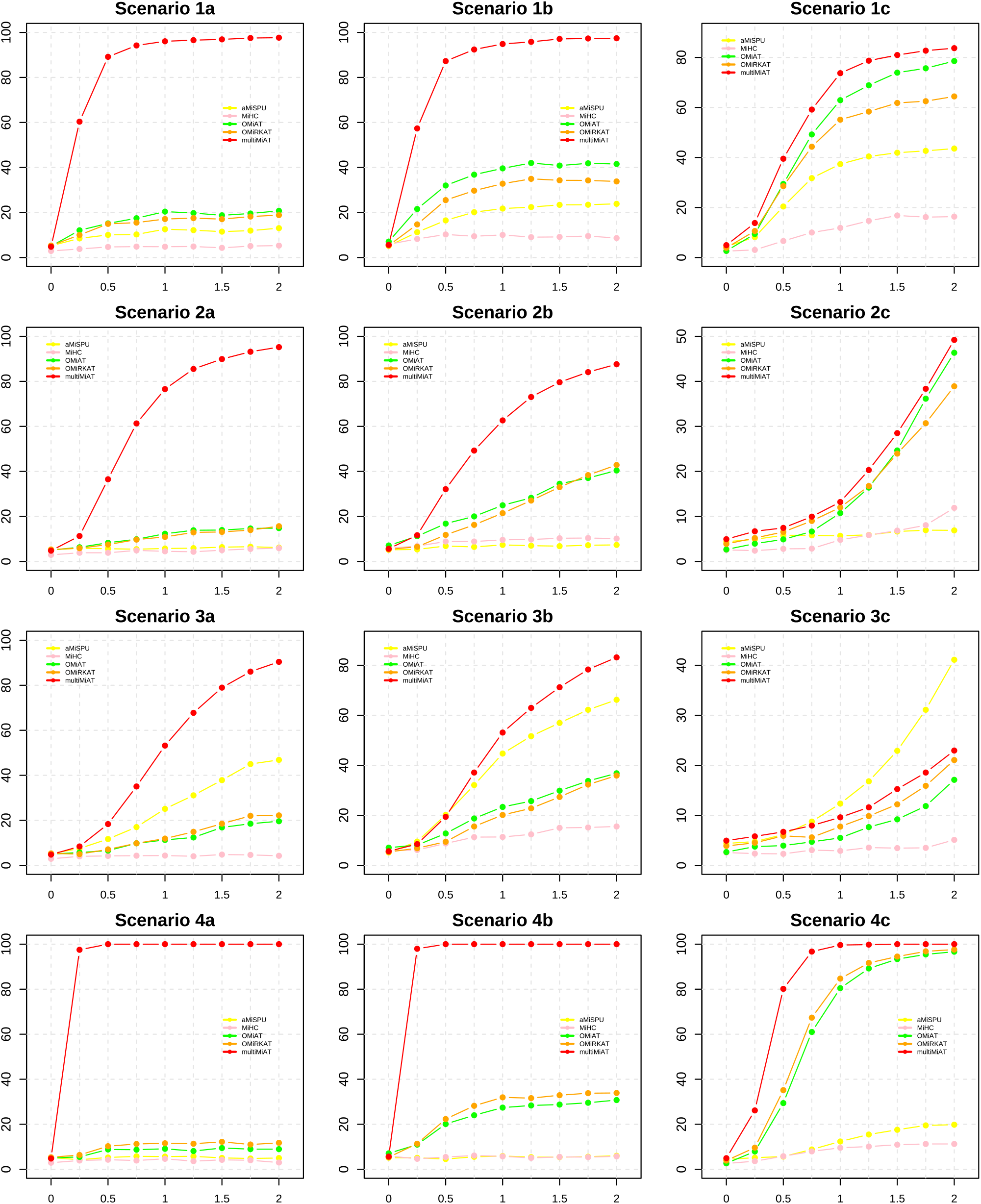
Under diverse scenarios, the comparison of the powers between our method (i.e., multiMiAT) and previous microbiome-based methods (*n* = 100).

**Figure 5.**
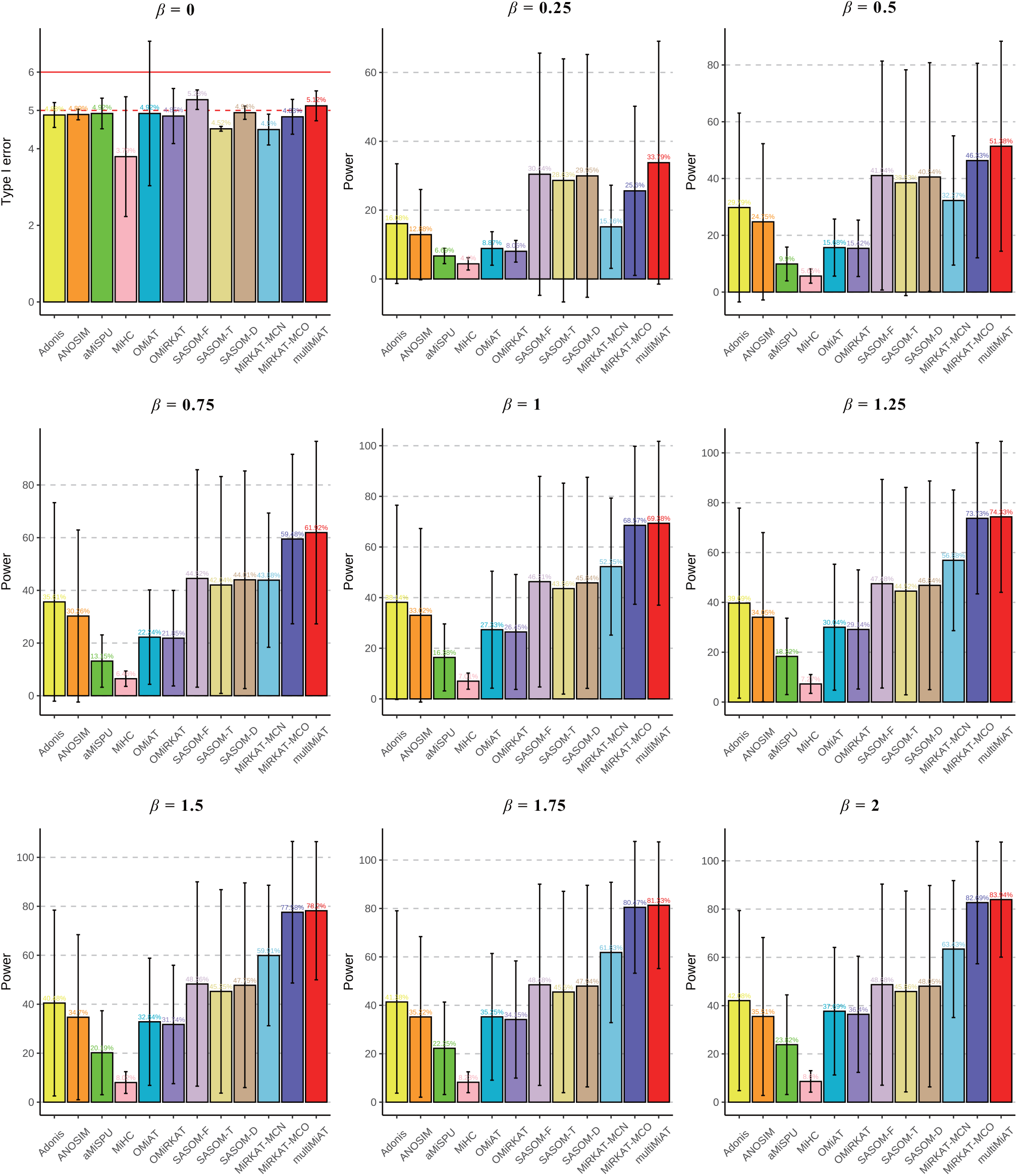
Comparison of empirical type I error rates and powers for synthesizing all scenarios under diverse effect sizes *β* (*n* = 100).

multiMiAT integrates three methods (i.e., multiMiRKAT, bcl-based score test and MiRKAT-MC), all of which have their own advantages in detecting different associated signals (Supplementary Figure S9, S10). Specifically, multiMiRKAT robustly provides competitive performance for nominal multicategory outcomes, while MiRKAT-MC is more advantageous in ordinal multicategory outcomes. In fact, also based on bcl, our method (i.e., multiMiRKAT-N) performs better than MiRKAT-MCN in some scenarios with ordinal outcomes (i.e., scenarios 1 and 4). In addition, the bcl-based score test has excellent performance in detecting abundant association signals. Overall, the optimal test multiMiAT is more powerful than these three methods, especially scenario 1, which further reflects their complementarity.

Compared with association tests for multicategory outcomes, our method robustly provides competitive performance for diverse scenarios (Figure 3, S16), especially abundant association signals (i.e., scenarios 1 and 4) and nominal multicategory outcomes (i.e., scenario c). Specifically, distance-based tests (i.e., Adonis and ANOSIM) do not take into account the influence of confounding factors, resulting in poor performance. In addition, the multicategory test SASOM is more sensitive to the situation where the association signals are relatively abundant (scenarios 1 and 4). Because of the lack of phylogenetic tree information, the effect is poor when the microbial association signal has a phylogenetic relationship (scenarios 1–3). MiRKAT-MCN and MiRKAT-MCO have advantages in detecting nonabundant association signals, especially rare association signals (scenario 3), but they are not powerful in detecting abundant association signals and nominal multicategory outcomes.

Compared with microbiome-based association tests for binary outcomes, we find that our methods are the best in all scenarios except scenario 3c (Figure 4, S17). Specifically, unbalanced outcomes have a certain negative impact on our method but a positive impact on microbiome-based association tests. Unbalanced outcomes increase the difference among adjacent groups, which makes the results of microbiome-based association tests more significant in adjacent pairwise analysis. aMiSPU has excellent performance in rare association signals (scenario 3) because aMiSPU contains unweighted individual tests that fully take the species presence/absence information into account.

To obtain a more convincing conclusion, we carried out a comprehensive analysis for all scenarios. We take the mean and standard deviation of all methods for these 12 scenarios. Although most methods varied significantly across different scenarios (large standard deviation), compared to all state-of-the-art methods, our methods consistently maintain the best performance (large mean, relatively stable variance) under diverse effect *β* sizes (Figure 5, S18). Specifically, when the effect size *β* = 0, all methods except OMiAT can control the type I error rate to approximately 5%. For *β* ≠ 0, multiMiAT has the best performance. With the increase in *β*, the advantage of our method compared with the other methods first decreases and then increases. In addition, when the sample size is small (i.e., *n* = 100), the advantage of our method is even more significant.

## Real data applications

In simulation experiments, we can control the association status between the microbiome and host phenotype through effect size; however, the true associations in real data experiments are unknown. Fortunately, a large number of studies have been developed to explore the association between the gut microbiome and disease, for example, colorectal cancer (CRC) [4, 15, 73–75] and *Clostridium difficile* infections (CDI) [76–78], which provides a reliable basis for our real data experiments. To fully verify the validity of our method, we adopt a small-scale dataset with balance outcomes (dataset A) and a large-scale real dataset with unbalance outcomes (dataset B).

### Association between the gut microbiome and colorectal cancer

To explore the characterization of the gut microbiome among three clinical statuses of CRC development (i.e., health, adenoma and carcinoma), Zackular et al. [73] collected stool samples from 90 subjects, where the number of healthy people, patients with colonic adenoma and patients with colonic adenocarcinoma were all 30. Age and race, the known clinical influencing factors of CRC, were considered for inclusion in our experiments. Here, race contains non-Hispanic whites and others. To filter out less abundant OTUs, we selected OTUs with the mean relative abundance ⩾ 10^−4^, which is a common preprocessing method [34, 43]. Through this preprocessing process, we obtained microbiome data containing 528 OTUs. We constructed a corresponding phylogenetic tree by using MEGA7 [79].

We find a relatively large difference among individual tests based on different dissimilarity measures. Specifically, all individual tests based on UniFrac distance can detect a significant association (i.e., *p* values less than 0.05); however, most individual tests based on other distances fail to reach the same conclusion (Supplementary Table S7), which is similar to the results of Adonis and ANOSIM analysis (Supplementary Figure S19). These results show that the gut microbiome associated with different statuses of CRC may vary widely in presence or absence. We summarized OTUs that were not present in all statuses, and we conducted species annotation using the function, *assignTaxonomy*, in the R package, *dada2* [80]. After filtering out OTUs that were not identified to the genus level, 17 OTUs are present in only one status, and 24 OTUs were present in two statuses (Supplementary Figure 20). Interestingly, many of these OTUs have been reported to be associated with CRC, for example, *Fusobacterium* [74, 75], *Bacteroides* [75]. Although the *p* values of Adonis and ANOSIM analysis based on UniFrac distance are less than 0.05, they are still higher than the *p* values of individual tests based on UniFrac distance. These methods ignore the influence of confounding factors, especially the impact of race on CRC. In fact, the proportion of whites was 84.4% in dataset A. However, the proportions of whites with different CRC statuses were 70.0% (health), 90.0% (adenoma) and 93.3% (carcinoma). In addition, most clm-based individual tests have better results than bcl-based individual tests with the same distance and kernel function control, which shows that there may be a progressive relationship among diverse statuses of CRC development.

Fortunately, our omnibus tests (i.e., multiMiRKAT-N and multiMiRKAT-O) and optimal test (multiMiAT) can also find a significant association (Supplementary Table S8). SASOM cannot find a significant association. Although MiRKAT-MC can find an association, the results of MiRKAT-MCN and MiRKAT-MCO are not more significant than those of multiMiRKAT-N and multiMiRKAT-O, respectively (Table 3, S8). Remarkably, any pair of statuses from the three statuses can be detected to be associated with the microbiome by at least one method of microbiome-based association tests (Table 3, S9). For pairwise, adjacent pairwise and baseline pairwise analyses of microbiome-based association tests, only some tests can detect significant associations (Table 3, S10), such as pairwise analysis of MiHC.

**Table 3.**
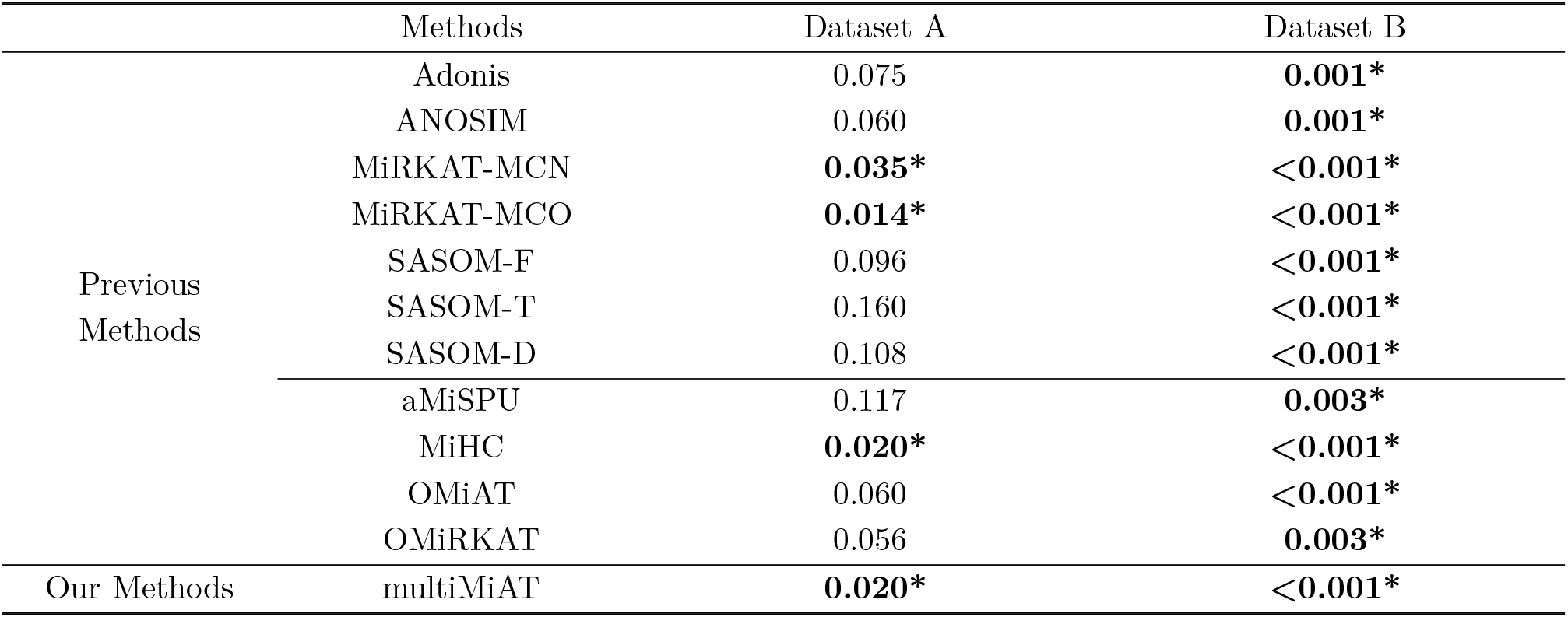
*P* values of our methods and previous methods in diverse datasets. Here, we conduct pairwise analysis of microbiome-based association test. ∗ represents a significant association (i.e., *p* value is below the significance level of 5%)

### Association between the gut microbiome and *Clostridium difficile* infections

To better understand the characterization of the gut microbiome among the diverse development statuses of CDI (i.e., nondiarrheal control, diarrheal control and case), Schubert et al. [76] collected fecal samples from 338 subjects. We selected age and gender as confounding factors. Age and gender were included in our real data analysis as potential confounding factors for CDI. We removed 3 samples with missing information on confounding factors from the dataset, where the preprocessed data contained 153 nondiarrheal controls, 89 diarrheal controls and 93 CDI patients. We also selected 439 OTUs with a mean relative abundance ⩾ 10^−4^ and constructed a corresponding phylogenetic tree by using MEGA7 [79].

Due to the large-scale real dataset, all our methods, including individual tests, omnibus tests and optimal tests, yield consistent conclusions: the extremely significant association between the gut microbiome and *Clostridium difficile* infections (Supplementary Table S7, S8). Unsurprisingly, all previous methods also detect this association (Table 3). In pairwise analysis of microbiome-based association tests, all tests have *p* values less than 0.05 (Supplementary Table S9, S10). In addition, the principal coordinate analyses (PCoA) based on diverse dissimilarities also reveal the differences in microbial composition among the diverse development statuses of CDI (Supplementary Figure S21).

## Discussion

In this paper, we propose a novel microbiome-based association test named multiMiAT to test the association between the microbiome and multicategory phenotypes. Most existing microbiome-based association tests are committed to detecting the association between the microbiome and continuous/binary phenotypes (i.e., BMI and disease), but fewer tests are designed for multicategory phenotypes. In addition, our simulation experiments demonstrate that our method can control the type I error rate correctly and conduct better performance in statistical power compared with state-of-the-art methods. Obviously, multiMiRKAT can be regarded as a tool to provide researchers with more complementary insights into multicategory phenotype studies.

In fact, the choices of multinomial logit model, dissimilarity measure and kernel function may have an important impact on the performance of individual tests in statistical power. We adopt multiple configurations simultaneously, such as constructing similarity metrics between subjects via different dissimilarity measures and kernel functions, to establish the corresponding individual tests. To maintain great statistical power in diverse scenarios, multiMiRKAT integrates these individual tests via *p* value combination methods. Here, the *p* value combination method does not simply take the best *p* value but further establishes the test statistic of the optimal test from these *p* values. Then, the final *p* value (i.e., *p* value of the optimal test) needs to be calculated according to the distribution of test statistics or the permutation method. Notably, the improper use of *p* value combination methods may not control type I error correctly or lose power. We verify the performance of three common combination methods and find that MinP is the most suitable compared with HMP and ACAT because the *p* values of individual tests based on diverse dissimilarity measures are greatly different in distribution. ACAT is sensitive, while HMP is conservative. Therefore, ACAT may obtain an inflated Type I error rate, and HMP may lose power.

Although the *p* value combination method is a popular strategy to maintain power for diverse scenarios, we find it difficult to guarantee that its results are always the best compared to all individual tests. The omnibus test usually produces noise accumulation during the combination process. Specifically, when most of the combined individual tests do not work well, it is difficult for the omnibus test to yield a satisfactory result unless other individual tests are sufficiently superior. Therefore, if the appropriate configuration is accurately obtained by learning some prior information, we can make the correct setting or optimize the configuration of the optimal test to greatly improve computational efficiency while ensuring higher power. In addition, unlike integrating individual tests with different configurations (i.e., dissimilarity measure, kernel function), an optimal test that combines different tests not only guarantees close to the best combined test but, in some cases, even outperforms the best combined test (Supplementary Figure S9, S10).

Our method still has much room for development. We find that our method does not work very well for detecting association signals with rare clusters (scenario 3) because these association signals may have difficulty reflecting the differences among different phenotypes in the similarity matrix. Other microbiome-based association tests for multicategory phenotypes can be considered for association signals with rare clusters, such as aMiSPU [32]. In addition, the microbiome-based association test for detecting the association between sparse association signals and multicategory phenotypes is also worth exploring, for example, the association between *Staphylococcus aureus* and disease severity of atopic dermatitis [81]. The optimal microbiome-based association test for multicategory phenotypes can also be extended to longitudinal data [47].

## Supporting information

Supplementary Material for "multiMiAT: An optimal microbiome-based test for multicategory phenotype"

## Acknowledgments

We would like to show our deepest gratitude to the studies of previous methods for giving us great inspiration, especially the MiRKAT-MC method. We acknowledge Biorender software (https://biorender.com/) which provides good picture materials.

## Author contributions statement

H.S. contributed to the methodological ideas for multiMiAT, performed the simulations and real data analyses, visualized the results, developed the software package, and wrote the manuscript. Y.W. contributed to the real data curation and analyses, visualized the results, and wrote the manuscript. Z.X. contributed to performed the simulations data analyses, and wrote the manuscript. X.H. visualized the results, and developed the software package. H.W. contributed to the real data curation and analyses, wrote the manuscript. T.H. contributed to the methodological ideas for multiMiAT, the biological insights and interpretations, and wrote the manuscript. X.J. contributed to the methodological ideas for multiMiAT, the biological insights and interpretations, and wrote the manuscript. All authors read and approved the final manuscript.

## Funding

This research is supported by the National Natural Science Foundation of China (61872157 and 61932008) and the Key Research and Development Program of Hubei Province (2020BAB017).

## Data availability

We used two public microbiome datasets in this paper: (1) the study on gut microbiome and colorectal cancer [73], whose all fastq files and clinical information are available at http://www.mothur.org/MicrobiomeBiomarkerCRC;(2) the study on gut microbiome and *Clostridium difficile* infections [76], whose all 16S rRNA gene sequence and clinical information are available at http://www.mothur.org/CDI_MicrobiomeModeling. These datasets can be also found at https://github.com/cduvallet/microbiomeHD [82]. All processed data including OTU count table, sample information and phylogenetic tree in our real data experiment is available at R package *multiMiAT*. Data and scripts used for all figures can be found at https://github.com/SunHan5/multiMiAT.SupplementaryMaterial.

## Notes

https://github.com/xpjiang-ccnu/multiMiAT

### Competing Interest Statement

The authors have declared no competing interest.

## References

1. Turnbaugh PJ, Ley RE, Hamady M, Fraser-Liggett CM, Knight R, and Gordon JI. The Human Microbiome Project. Nature, 449(7164):804–810, 2007.

2. Koeth RA, Wang Z, Levison BS, Buffa JA, Org E, Sheehy BT, Britt EB, Fu X, Wu Y, Li L, and et al. Intestinal microbiota metabolism of L-carnitine, a nutrient in red meat, promotes atherosclerosis. Nat Med, 19(5):576–585, 2013.

3. Zitvogel L, Ma Y, Raoult D, Kroemer G, and Gajewski TF. The microbiome in cancer immunotherapy: Diagnostic tools and therapeutic strategies. Science, 359(6382):1366–1370, 2018.

4. Pushalkar S, Hundeyin M, Daley D, Zambirinis CP, Kurz E, Mishra A, Mohan N, Aykut B, Usyk M, Torres LE, and et al. The Pancreatic Cancer Microbiome Promotes Oncogenesis by Induction of Innate and Adaptive Immune Suppression. Cancer Discov, 8(4):403–416, 2018.

5. Helmink BA, Khan MAW, Hermann A, Gopalakrishnan V, and Wargo JA. The microbiome, cancer, and cancer therapy. Nat Med, 25(3):377–388, 2019.

6. Sommer F and Bäckhed F. The gut microbiota – masters of host development and physiology. Nat Rev Microbiol, 11(4):227–238, 2013.

7. Sender R, Fuchs S, and Milo R. Are We Really Vastly Outnumbered? Revisiting the Ratio of Bacterial to Host Cells in Humans. Cell, 164(3):337–340, 2016.

8. Gilbert JA, Blaser MJ, Caporaso JG, Jansson JK, Lynch SV, and Knight R. Current understanding of the human microbiome. Nat Med, 24(4):392–400, 2018.

9. Ventura M, O’Flaherty S, Claesson MJ, Turroni F, Klaenhammer TR, van Sinderen D, and O’Toole PW. Genome-scale analyses of health-promoting bacteria: probiogenomics. Nat Rev Microbiol, 7(1):61–71, 2009.

10. Hsiao EY, McBride SW, Hsien S, Sharon G, Hyde ER, McCue T, Codelli JA, Chow J, Reisman SE, Petrosino JF, and et al. Microbiota Modulate Behavioral and Physiological Abnormalities Associated with Neurodevelopmental Disorders. Cell, 155(7):1451–1463, 2013.

11. Sanna S, van Zuydam NR, Mahajan A, Kurilshikov A, Vich Vila A, Võsa U, Mujagic Z, Masclee AAM, Jonkers DMAE, Oosting M, Joosten LAB, and et al. Causal relationships among the gut microbiome, short-chain fatty acids and metabolic diseases. Nat Genet, 51(4):600–605, 2019.

12. Jensen BAH and Marette A. Microbial translocation in type 2 diabetes: when bacterial invaders overcome host defence in human obesity. Gut, 69(10):1724–1726, 2020.

13. Mayhew D, Devos N, Lambert C, Brown JR, Clarke SC, Kim VL, Magid-Slav M, Miller BE, Ostridge KK, Patel R, and et al. Longitudinal profiling of the lung microbiome in the AERIS study demonstrates repeatability of bacterial and eosinophilic COPD exacerbations. Thorax, 73(5):422–430, 2018.

14. Ditz B, Christenson S, Rossen J, Brightling C, Kerstjens HAM, van den Berge M, and Faiz A. Sputum microbiome profiling in COPD: beyond singular pathogen detection. Thorax, 75(4):338–344, 2020.

15. Hale VL, Jeraldo P, Chen J, Mundy M, Yao J, Priya S, Keeney G, Lyke K, Ridlon J, White BA, and et al. Distinct microbes, metabolites, and ecologies define the microbiome in deficient and proficient mismatch repair colorectal cancers. Genome Med, 10(1):78, 2018.

16. Gehrig JL, Venkatesh S, Chang H, Hibberd MC, Kung VL, Cheng J, Chen RY, Subramanian S, Cowardin CA, Meier MF, and et al. Effects of microbiota-directed foods in gnotobiotic animals and undernourished children. Science, 365(6449):eaau4732, 2019.

17. Raman AS, Gehrig JL, Venkatesh S, Chang H, Hibberd MC, Subramanian S, Kang G, Bessong PO, Lima AAM, Kosek MN, and et al. A sparse covarying unit that describes healthy and impaired human gut microbiota development. Science, 365(6449):eaau4735, 2019.

18. Bashir A, Klammer AA, Robins WP, Chin C, Webster D, Paxinos E, Hsu D, Ashby M, Wang S, Peluso P, and et al. A hybrid approach for the automated finishing of bacterial genomes. Nat Biotechnol, 30(7):701–707, 2012.

19. Liu Y, Qin Y, Chen T, Lu M, Qian X, Guo X, and Bai Y. A practical guide to amplicon and metagenomic analysis of microbiome data. Protein Cell, 12(5):315–330, 2021.

20. Caporaso JG, Kuczynski J, Stombaugh J, Bittinger K, Bushman FD, Costello EK, Fierer N, Peña AG, Goodrich JK, Gordon JI, and et al. QIIME allows analysis of high-throughput community sequencing data. Nat Methods, 7(5):335–336, 2010.

21. Hamady M and Knight R. Microbial community profiling for human microbiome projects: Tools, techniques, and challenges. Genome Research, 19(7):1141–1152, 2009.

22. Price ND, Magis AT, Earls JC, Glusman G, Levy R, Lausted C, McDonald DT, Kusebauch U, Moss CL, Zhou Y, and et al. A wellness study of 108 individuals using personal, dense, dynamic data clouds. Nat Biotechnol, 35(8):747–756, 2017.

23. Anderson MJ. A new method for non-parametric multivariate analysis of variance. Austral Ecology, 26(1):32–46, 2001.

24. Clarke KR. Non-parametric multivariate analyses of changes in community structure. Austral Ecol, 18(1):117–143, 1993.

25. Warton DI, Wright ST, and Wang Y. Distance-based multivariate analyses confound location and dispersion effects. Methods in Ecology and Evolution, 3(1):89–101, 2012.

26. Mars RAT, Yang Y, Ward T, Houtti M, Priya S, Lekatz HR, Tang X, Sun Z, Kalari KR, Korem T, and et al. Longitudinal Multi-omics Reveals Subset-Specific Mechanisms Underlying Irritable Bowel Syndrome. Cell, 182(6):1460–1473, 2020.

27. Bramble MS, Vashist N, Ko A, Priya S, Musasa C, Mathieu A, Spencer DA, Lupamba Kasendue M, Mamona Dilufwasayo P, Karume K, and et al. The gut microbiome in konzo. Nat Commun, 12(1):5371, 2021.

28. Schirmer M, Denson L, Vlamakis H, Franzosa EA, Thomas S, Gotman NM, Rufo P, Baker SS, Sauer C, Markowitz J, and et al. Compositional and Temporal Changes in the Gut Microbiome of Pediatric Ulcerative Colitis Patients Are Linked to Disease Course. Cell Host & Microbe, 24(4):600–610, 2018.

29. Brand EC, Klaassen MAY, Gacesa R, Vich Vila A, Ghosh H, de Zoete MR, Boomsma DI, Hoentjen F, Horjus Talabur Horje CS, van de Meeberg PC, and et al. Healthy Cotwins Share Gut Microbiome Signatures With Their Inflammatory Bowel Disease Twins and Unrelated Patients. Gastroenterology, 160(6):1970–1985, 2021.

30. Tang Z, Chen G, and Alekseyenko AV. PERMANOVA-S: association test for microbial community composition that accommodates confounders and multiple distances. Bioinformatics, 32(17):2618–2625, 2016.

31. Zhao N, Chen J, Carroll IM, Ringel-Kulka T, Epstein MP, Zhou H, Zhou JJ, Ringel Y, Li H, and Wu MC. Testing in Microbiome-Profiling Studies with MiRKAT, the Microbiome Regression-Based Kernel Association Test. The American Journal of Human Genetics, 96(5):797–807, 2015.

32. Wu C, Chen J, Kim J, and Pan W. An adaptive association test for microbiome data. Genome Med, 8(1):56, 2016.

33. Koh H. An adaptive microbiome α-diversity-based association analysis method. Sci Rep, 8(1):18026, 2018.

34. Koh H and Zhao N. A powerful microbial group association test based on the higher criticism analysis for sparse microbial association signals. Microbiome, 8(1):63, 2020.

35. Wilson N, Zhao N, Zhan X, Koh H, Fu W, Chen J, Li H, Wu MC, and Plantinga AM. MiRKAT: kernel machine regression-based global association tests for the microbiome. Bioinformatics, 37(11):1595–1597, 2021.

36. Plantinga A, Zhan X, Zhao N, Chen J, Jenq RR, and Wu MC. MiRKAT-S: a community-level test of association between the microbiota and survival times. Microbiome, 5(1):17, 2017.

37. Koh H, Livanos AE, Blaser MJ, and Li H. A highly adaptive microbiome-based association test for survival traits. BMC Genomics, 19(1):210, 2018.

38. Zhan X, Xue L, Zheng H, Plantinga A, Wu MC, Schaid DJ, Zhao N, and Chen J. A small-sample kernel association test for correlated data with application to microbiome association studies. Genet. Epidemiol., 42(8):772–782, 2018.

39. Koh H, Li Y, Zhan X, Chen J, and Zhao N. A Distance-Based Kernel Association Test Based on the Generalized Linear Mixed Model for Correlated Microbiome Studies. Front. Genet., 10:458, 2019.

40. Sun H, Huang X, Fu L, Huo B, He T, and Jiang X. A powerful adaptive microbiome-based association test for microbial association signals with diverse sparsity levels. Journal of Genetics and Genomics, 48(9):851–859, 2021.

41. Zhan X, Tong X, Zhao N, Maity A, Wu MC, and Chen J. A small-sample multivariate kernel machine test for microbiome association studies. Genet. Epidemiol., 41(3):210–220, 2017.

42. Zhan X, Plantinga A, Zhao N, and Wu MC. A fast small-sample kernel independence test for microbiome community-level association analysis. Biom, 73(4):1453–1463, 2017.

43. Koh H, Blaser MJ, and Li H. A powerful microbiome-based association test and a microbial taxa discovery framework for comprehensive association mapping. Microbiome, 5(1):45, 2017.

44. Wang T, Ling W, Plantinga AM, Wu MC, and Zhan X. Testing microbiome association using integrated quantile regression models. Bioinformatics, 38(2):419–425, 2022.

45. Banerjee K, Chen J, and Zhan X. Adaptive and powerful microbiome multivariate association analysis via feature selection. NAR Genomics and Bioinformatics, 4(1):lqab120, 2022.

46. Zhang L, Wang Y, Chen J, and Chen J. RFtest: A Robust and Flexible Community-Level Test for Microbiome Data Powerfully Detects Phylogenetically Clustered Signals. Frontiers in Genetics, 12:749573, 2022.

47. Jiang Z, He M, Chen J, Zhao N, and Zhan X. MiRKAT-MC: A Distance-Based Microbiome Kernel Association Test With Multi-Categorical Outcomes. Frontiers in Genetics, 13:841764, 2022.

48. Sun H, Huang X, Huo B, Tan Y, He T, and Jiang X. Detecting sparse microbial association signals adaptively from longitudinal microbiome data based on generalized estimating equations. Briefings in Bioinformatics, page bbac149, 2022.

49. Touloumis A, Agresti A, and Kateri M. GEE for Multinomial Responses Using a Local Odds Ratios Parameterization: GEE for Multinomial Responses Using a Local Odds Ratios Parameterization. Biom, 69(3):633–640, 2013.

50. David LA, Maurice CF, Carmody RN, Gootenberg DB, Button JE, Wolfe BE, Ling AV, Devlin AS, Varma Y, Fischbach MA, and et al. Diet rapidly and reproducibly alters the human gut microbiome. Nature, 505(7484):559–563, 2014.

51. Bi W, Zhou W, Dey R, Mukherjee B, Sampson JN, and Lee S. Efficient mixed model approach for largescale genome-wide association studies of ordinal categorical phenotypes. The American Journal of Human Genetics, 108(5):825–839, 2021.

52. Li Y, Ke Y, Xia X, Wang Y, Cheng F, Liu X, Jin X, Li B, Xie C, Liu S, and et al. Genome-wide association study of COVID-19 severity among the Chinese population. Cell Discov, 7(1):76, 2021.

53. Liu H, Chen X, Hu X, Niu H, Tian R, Wang H, Pang H, Jiang L, Qiu B, Chen X, and et al. Alterations in the gut microbiome and metabolism with coronary artery disease severity. Microbiome, 7(1):68, 2019.

54. Jha AR, Davenport ER, Gautam Y, Bhandari D, Tandukar S, Ng KM, Fragiadakis GK, Holmes S, Gautam GP, Leach J, and et al. Gut microbiome transition across a lifestyle gradient in Himalaya. PLoS Biol, 16(11):e2005396, 2018.

55. Liu M, Liu Y, Wu MC, Hsu L, and He Q. A method for subtype analysis with somatic mutations. Bioinformatics, 37(1):50–56, 2021.

56. Maity A, Sullivan PF, and Tzeng J. Multivariate Phenotype Association Analysis by Marker-Set Kernel Machine Regression: Multivariate Kernel Machine Regression. Genet. Epidemiol., 36(7):686–695, 2012.

57. Agresti A. Categorical data analysis, 3rd ed. Wiley, Hoboken, NJ, 2013.

58. Agarwal D and Zhang NR. Semblance: An empirical similarity kernel on probability spaces. Science Advances, 5(12):eaau9630, 2019.

59. Touloumis A. R Package multgee : A Generalized Estimating Equations Solver for Multinomial Responses. J. Stat. Soft., 64(8), 2015.

60. Bray JR and Curtis JT. An Ordination of the Upland Forest Communities of Southern Wisconsin. Ecological Monographs, 27(4):325–349, 1957.

61. Lozupone C and Knight R. UniFrac: a New Phylogenetic Method for Comparing Microbial Communities. Appl Environ Microbiol, 71(12):8228–8235, 2005.

62. Lozupone CA, Hamady M, Kelley ST, and Knight R. Quantitative and Qualitative Diversity Measures Lead to Different Insights into Factors That Structure Microbial Communities. Appl Environ Microbiol, 73(5):1576–1585, 2007.

63. Chen J, Bittinger K, Charlson ES, Hoffmann C, Lewis J, Wu GD, Collman RG, Bushman FD, and Li H. Associating microbiome composition with environmental covariates using generalized UniFrac distances. Bioinformatics, 28(16):2106–2113, 2012.

64. Jaccard P. THE DISTRIBUTION OF THE FLORA IN THE ALPINE ZONE. New Phytol, 11(2):37–50, 1912.

65. Liu Y, Chen S, Li Z, Morrison AC, Boerwinkle E, and Lin X. ACAT: A Fast and Powerful p Value Combination Method for Rare-Variant Analysis in Sequencing Studies. The American Journal of Human Genetics, 104(3):410–421, 2019.

66. Wilson DJ. The harmonic mean p-value for combining dependent tests. Proceedings of the National Academy of Sciences, 116(4):1195–1200, 2019.

67. Schaid DJ, Rowland CM, Tines DE, Jacobson RM, and Poland GA. Score Tests for Association between Traits and Haplotypes when Linkage Phase Is Ambiguous. The American Journal of Human Genetics, 70(2):425–434, 2002.

68. Kim J, Zhang Y, and Pan W. Powerful and Adaptive Testing for Multi-trait and Multi-SNP Associations with GWAS and Sequencing Data. Genetics, 203(2):715–731, 2016.

69. He Q, Liu Y, Liu M, Wu MC, and Hsu L. Random effect based tests for multinomial logistic regression in genetic association studies. Genetic Epidemiology, 45(7):736–740, 2021.

70. Touloumis A. Simulating Correlated Binary and Multinomial Responses under Marginal Model Specification: The SimCorMultRes Package. The R Journal, 8(2):79, 2016.

71. Charlson ES, Chen J, Custers-Allen R, Bittinger K, Li H, Sinha R, Hwang J, Bushman FD, and Collman RG. Disordered Microbial Communities in the Upper Respiratory Tract of Cigarette Smokers. PLoS ONE, 5(12):e15216, 2010.

72. Reynolds AP, Richards G, de la Iglesia B, and Rayward-Smith VJ. Clustering Rules: A Comparison of Partitioning and Hierarchical Clustering Algorithms. J Math Model Algor, 5(4):475–504, 2006.

73. Zackular JP, Rogers MAM, Ruffin MT, and Schloss PD. The Human Gut Microbiome as a Screening Tool for Colorectal Cancer. Cancer Prevention Research, 7(11):1112–1121, 2014.

74. Zhou Z, Chen J, Yao H, and Hu H. Fusobacterium and Colorectal Cancer. Frontiers in Oncology, 8:371, 2018.

75. Zagato E, Pozzi C, Bertocchi A, Schioppa T, Saccheri F, Guglietta S, Fosso B, Melocchi L, Nizzoli G, Troisi J, and et al. Endogenous murine microbiota member Faecalibaculum rodentium and its human homologue protect from intestinal tumour growth. Nature Microbiology, 5(3):511–524, 2020.

76. Schubert AM, Rogers MAM, Ring C, Mogle J, Petrosino JP, Young VB, Aronoff DM, and Schloss PD. Microbiome Data Distinguish Patients with Clostridium difficile Infection and Non-C. difficile-Associated Diarrhea from Healthy Controls. mBio, 5(3):e01021–14, 2014.

77. Zackular JP, Moore JL, Jordan AT, Juttukonda LJ, Noto MJ, Nicholson MR, Crews JD, Semler MW, Zhang Y, Ware LB, and et al. Dietary zinc alters the microbiota and decreases resistance to Clostridium difficile infection. Nature Medicine, 22(11):1330–1334, 2016.

78. Nagao-Kitamoto H, Leslie JL, Kitamoto S, Jin C, Thomsson KA, Gillilland MG, Kuffa P, Goto Y, Jenq RR, Ishii C, and et al. Interleukin-22-mediated host glycosylation prevents Clostridioides difficile infection by modulating the metabolic activity of the gut microbiota. Nature Medicine, 26(4):608–617, 2020.

79. Kumar S, Stecher G, and Tamura K. MEGA7: Molecular Evolutionary Genetics Analysis Version 7.0 for Bigger Datasets. Molecular Biology and Evolution, 33(7):1870–1874, 2016.

80. Callahan BJ, McMurdie PJ, Rosen MJ, Han AW, Johnson AJA, and Holmes SP. DADA2: High-resolution sample inference from Illumina amplicon data. Nature Methods, 13(7):581–583, 2016.

81. Kong HH, Oh J, Deming C, Conlan S, Grice EA, Beatson MA, Nomicos E, Polley EC, Komarow HD, NISC Comparative Sequence Program, and et al. Temporal shifts in the skin microbiome associated with disease flares and treatment in children with atopic dermatitis. Genome Research, 22(5):850–859, 2012.

82. Duvallet C, Gibbons SM, Gurry T, Irizarry RA, and Alm EJ. Meta-analysis of gut microbiome studies identifies disease-specific and shared responses. Nature Communications, 8(1):1784, 2017.

