## Supplementary Material for "multiMiAT: An optimal microbiome-based test for multicategory phenotype" for "multiMiAT: An optimal microbiome-based association test for multicategory phenotypes"

### 1 Calculation process for the $p$ value of multiMiAT test

We calculate $p$ value via the following steps:

1. **Residuals from the fitted multinomial logit models:** The null models (i.e., bcl and clm) are fitted and the residuals $\hat{\mathbf{r}}_{l(0)}$ are estimated via $\hat{\mathbf{r}} = (y_{11} - \hat{\mu}_{11}, \ldots, y_{n(J-1)} - \hat{\mu}_{n(J-1)})^T$ [1], where $l \in \Gamma_{\mathrm{L}} = \{\mathrm{M_{bcl}}, \mathrm{M_{clm}}\}$. $\hat{\mu}_{ij}$ is the expectation under the null model, where $i = 1, 2, \ldots, n$, $j = 1, 2, \ldots, J$, $J > 2$. We can obtain the $b$th permutated residual $\hat{\mathbf{r}}_{l(b)}$ by reshuffling the residuals $\hat{\mathbf{r}}_{l(0)}$, where $b \in \{1, 2, \ldots, B\}$.

2. **Similarity matrix between subjects:** We calculate the dissimilarity matrices between subjects according to difference dissimilarity measures $\Gamma_{\mathrm{D}} = \{\mathrm{D_{BC}}, \mathrm{D_u}, \mathrm{D_0}, \mathrm{D_{0.25}}, \mathrm{D_{0.5}}, \mathrm{D_{0.75}}, \mathrm{D_w}\}$ [2, 3, 4, 5]. Then, the similarity matrices are obtained by transforming the dissimilarity matrix between subjects through diverse kernel functions $\Gamma_{\mathrm{K}} = \{\mathrm{K_{LK}}, \mathrm{K_{GK}}, \mathrm{K_{LaK}}\}$ [6]. The kernel functions contain

   1) Linear kernel ($\mathrm{K_{LK}}$) function: $\mathbf{K}_d = -\frac{1}{2}(\mathbf{I}_n - \frac{\mathbf{1}_n\mathbf{1}_n^T}{n})\mathbf{D}_d^2(\mathbf{I}_n - \frac{\mathbf{1}_n\mathbf{1}_n^T}{n})$,
   2) Gaussian kernel ($\mathrm{K_{GK}}$) function: $\mathbf{K}_d = \exp(-\frac{\mathbf{D}_d^2}{2\sigma^2})$,
   3) Laplacian kernel ($\mathrm{K_{LaK}}$) function: $\mathbf{K}_d = \exp(-\frac{\mathbf{D}_d}{\sigma})$,

   where $\mathbf{I}_n$ is a $n \times n$ identity matrix, $\mathbf{1}_n$ is a $n$-dimensional vector of ones. $\mathbf{D}_d$ is dissimilarity matrix and $d \in \Gamma_{\mathrm{D}}$. $\sigma$ is the hyperparameter.

3. **Test statistics and $p$ values of microbiome regression-based kernel individual tests:** The reconstructed kernel matrix $\widetilde{\mathbf{K}}_{kd}$ are calculated according to kernel function and eigenvalue decomposition [7]. The null test statistic $T_l^{kd}$ and permutated test statistic $T_{l(b)}^{kd}$ can be obtained via

   $$T_l^{kd} = \hat{\mathbf{r}}_l^T \mathbf{V}_l^{-1} \widetilde{\mathbf{K}}_{kd} \mathbf{V}_l^{-1} \hat{\mathbf{r}}_l,$$

   where $\mathbf{V}_l = \mathbf{I}_n \otimes \mathbf{V}_{l0}$ and $\mathbf{V}_{l0}$ is the estimated residual variance matrix under the null model. Then, the $p$ value $p_l^{kd}$ can be calculated through

   $$p_l^{kd} = [\sum_{b=1}^{B} I(T_{l(b)}^{kd} > T_l^{kd}) + 0.01]/(B + 0.01).$$

4. **Test statistics and $p$ values of microbiome regression-based kernel omnibus tests multiMiRKAT (i.e., multiMiRKAT-N and multiMiRKAT-O):** We first calculate $p_{l(b)}^{kd}$ via

   $$p_{l(b)}^{kd} = [\sum_{b' \neq b} I(T_{l(b')}^{kd} > T_{l(b)}^{kd}) + 0.01]/(B + 0.01).$$

   Then, the test statistics $T_{\mathrm{multiMiRKAT}}$ and permutated test statistic $T_{\mathrm{multiMiRKAT}(b)}$ based on bcl and clm are as follows

   $$T_{\mathrm{multiMiRKAT}} = \min_{k \in \Gamma_{\mathrm{K}}} \min_{d \in \Gamma_{\mathrm{D}}} p_l^{kd},$$
   $$T_{\mathrm{multiMiRKAT}(b)} = \min_{k \in \Gamma_{\mathrm{K}}} \min_{d \in \Gamma_{\mathrm{D}}} p_{l(b)}^{kd}.$$

   And the $p$ values $p_{\mathrm{multiMiRKAT}}$ can be obtained via

   $$p_{\mathrm{multiMiRKAT}} = [\sum_{b=1}^{B} I(T_{\mathrm{multiMiRKAT}(b)} < T_{\mathrm{multiMiRKAT}}) + 0.01]/(B + 0.01).$$

5. **Test statistics and $p$ value of score test [8]:** We first calculate the covariance matrix $\mathbf{\Sigma}$ of multinomial outcomes $\overline{\mathbf{Y}}$. Then we can obtain $\mathbf{U} = \mathbf{S}^T(\overline{\mathbf{Y}} - \overline{\boldsymbol{\mu}})$ and the covariance matrix $\widehat{\mathbf{\Sigma}}$ of $\mathbf{U}$, where $\mathbf{S} = \mathbf{I}_{(J-1)} \otimes (\mathbf{OW}^T)$ and $\widehat{\mathbf{\Sigma}} = \mathbf{S}^T(\mathbf{\Sigma} - \mathbf{\Sigma}\mathbb{X}(\mathbb{X}^T\mathbf{\Sigma}\mathbb{X})^{-1}\mathbb{X}^T\mathbf{\Sigma})\mathbf{S})$. The test statistics $T_{\mathrm{score}}$ of score test can be established through

   $$T_{\mathrm{score}} = \mathbf{U}^T \widehat{\mathbf{\Sigma}}^{-1} \mathbf{U}.$$

   Because $T_{\mathrm{score}}$ can be shown to asymptotically follow a $\chi^2$ distribution with degree of freedom $(J - 1)$, we can obtain the $p$ value of score test $p_{\mathrm{score}}$ from asymptotic distribution.

6. **Test statistics and $p$ values of MiRKAT-MC** [9]**:** The test statistics of individual tests $Q_l^{kd}$ can be obtained through

$$Q_l^{kd} = (\overline{\mathbf{Y}} - \overline{\boldsymbol{\mu}})^T \overline{\mathbf{D}}^T \overline{\mathbf{W}}\, \overline{\mathbf{K}}_{kd} \overline{\mathbf{W}}\, \overline{\mathbf{D}} (\overline{\mathbf{Y}} - \overline{\boldsymbol{\mu}}),$$

where $\overline{\mathbf{K}}_{kd} = \mathbf{I}_{J-1} \otimes \mathbf{K}_{kd}^*$. $\overline{\mathbf{D}} = \partial(\mathbb{X}\hat{\boldsymbol{\beta}})/\partial\boldsymbol{\mu}$ and $\overline{\mathbf{W}} = (\overline{\mathbf{D}}\,\overline{\mathbf{V}}\,\overline{\mathbf{D}})^{-1}$. Then, we can obtain the $p$ values of individual tests via pseudo-permutation strategy, that is, all permuted test statistic can be calculated based on the Pearson type III density [10]. And MiRKAT-MC test uses the HMP [11] to combine these $p$ values of individual tests, where MiRKAT-MCN test and MiRKAT-MCO test are bcl-based and clm-based MiRKAT-MC test, respectively.

7. **Test statistics and $p$ values of optimal tests multiMiAT:** Considering that the $p$ value of score test is obtained based on asymptotic distribution rather than permutation method, we establish the test statistics $T_{\text{multiMiAT}}$ of multiMiAT test through HMP [11], whose model is as follows

$$T_{\text{multiMiAT}} = \frac{\sum_{t \in \Gamma_{\mathrm{T}}} w_t}{\sum_{t \in \Gamma_{\mathrm{T}}} w_t / p_t},$$

where $\sum_{t \in \Gamma_{\mathrm{T}}} w_t = 1$ and $\Gamma_{\mathrm{T}} = \{T_{\text{multiMiRKAT-N}}, T_{\text{multiMiRKAT-O}}, T_{\text{score}}, T_{\text{MiRKAT-MCN}}, T_{\text{MiRKAT-MCO}}\}$. $p_t$ represents the $p$ values of these combined tests. And the $p$ values $p_{\text{multiMiAT}}$ can be obtained.

$$p_{\text{multiMiAT}} = \int_{1/T_{\text{multiMiAT}}}^{\infty} f_{\text{Landau}}(x|\log L + c, \frac{\pi}{2})\mathrm{d}x,$$

where $f_{\text{Landau}}(x|\mu, \sigma) = \frac{1}{\pi\sigma}\int_0^\infty e^{-t\frac{(x-\mu)}{\sigma} - \frac{2}{\pi} t \log t} \sin(2t)\mathrm{d}t$ and $c = 1 + \psi(1) - \log(2/\pi) \approx 0.874$. $L$ is the number of combined $p$ values (i.e., $|\Gamma_{\mathrm{T}}| = 5$) and $\psi(x) = \mathrm{d}/\mathrm{d}x\{\ln\mathit{\Gamma}(x)\} = \mathit{\Gamma}'(x)/\mathit{\Gamma}(x)$.

### 2 Supplementary Tables

#### 2.1 Analysis for type I error in simulation experiment

Table S1: Empirical type I error rates of microbiome regression-based kernel local omnibus tests which use $p$ value combination method (MinP) to integrate the individual tests. Here, these individual tests are with diverse dissimilarity measures.

| Model | Kernel | $n = 100$ | | | $n = 200$ | | |
|---|---|---|---|---|---|---|---|
| | | Ordinal | | Nominal | Ordinal | | Nominal |
| | | Balance | Unbalance | | Balance | Unbalance | |
| bcl | LK | 5.24% | 5.12% | 5.32% | 4.68% | 5.42% | 5.28% |
| | GK | 5.38% | 5.06% | 5.42% | 4.62% | 5.52% | 5.38% |
| | LaK | 5.20% | 5.04% | 5.16% | 4.64% | 5.30% | 5.26% |
| clm | LK | 5.12% | 4.94% | 5.14% | 4.88% | 5.18% | 4.72% |
| | GK | 5.12% | 4.86% | 5.10% | 4.76% | 5.08% | 4.84% |
| | LaK | 5.26% | 5.00% | 5.02% | 4.60% | 5.10% | 4.80% |

Table S2: Empirical type I error rates of microbiome regression-based kernel local omnibus tests which use $p$ value combination method (HMP) to integrate the individual tests. Here, these individual tests are with diverse dissimilarity measures.

| Model | Kernel | $n = 100$ | | | $n = 200$ | | |
|---|---|---|---|---|---|---|---|
| | | Ordinal | | Nominal | Ordinal | | Nominal |
| | | Balance | Unbalance | | Balance | Unbalance | |
| bcl | LK | 4.98% | 4.60% | 4.72% | 4.26% | 4.96% | 4.92% |
| | GK | 5.00% | 4.64% | 4.66% | 4.18% | 5.08% | 4.90% |
| | LaK | 4.82% | 4.70% | 4.80% | 4.16% | 4.90% | 4.82% |
| clm | LK | 5.00% | 4.50% | 4.96% | 4.24% | 4.86% | 4.56% |
| | GK | 4.86% | 4.46% | 4.88% | 4.26% | 4.90% | 4.62% |
| | LaK | 4.86% | 4.54% | 4.90% | 4.40% | 4.78% | 4.68% |

Table S3: Empirical type I error rates of microbiome regression-based kernel local omnibus tests which use $p$ value combination method (ACAT [12]) to integrate the individual tests. Here, these individual tests are with diverse dissimilarity measures.

| Model | Kernel | $n$ = 100 | | | $n$ = 200 | | |
|---|---|---|---|---|---|---|---|
| | | Ordinal | | Nominal | Ordinal | | Nominal |
| | | Balance | Unbalance | | Balance | Unbalance | |
| bcl | LK | 6.12%* | 5.74% | 6.04%* | 5.36% | 6.14%* | 6.06%* |
| | GK | 6.08%* | 5.80% | 6.00%* | 5.44% | 6.18%* | 6.10%* |
| | LaK | 6.00%* | 5.64% | 6.02%* | 5.52% | 6.10%* | 6.08%* |
| clm | LK | 6.08%* | 5.68% | 6.12%* | 5.52% | 6.06%* | 6.00%* |
| | GK | 6.10%* | 5.74% | 6.06%* | 5.40% | 5.94% | 5.90% |
| | LaK | 6.10%* | 5.66% | 6.20%* | 5.44% | 5.96% | 5.78% |

Table S4: Empirical type I error rates of score tests, microbiome regression-based kernel global omnibus tests (i.e., multiMiRKAT-N and multiMiRKAT-O) and optimal tests (i.e., multiMiAT). Our method multiMiAT uses MinP and HMP for $p$ value combination. * represents inflated type I error rates.

| Category | Method | $n$ = 100 | | | $n$ = 200 | | |
|---|---|---|---|---|---|---|---|
| | | Ordinal | | Nominal | Ordinal | | Nominal |
| | | Balance | Unbalance | | Balance | Unbalance | |
| Individual test | Score test | 5.60% | 5.62% | 5.04% | 5.14% | 4.98% | 4.62% |
| Omnibus test | multiMiRKAT-N(MinP) | 5.28% | 5.08% | 5.20% | 4.66% | 5.46% | 5.42% |
| | multiMiRKAT-N(HMP) | 5.10% | 4.76% | 4.82% | 4.26% | 5.06% | 4.96% |
| | multiMiRKAT-N(ACAT) | 6.08%* | 5.78% | 5.94% | 5.54% | 6.08%* | 6.22%* |
| | multiMiRKAT-O(MinP) | 5.30% | 4.94% | 5.08% | 4.80% | 5.06% | 4.80% |
| | multiMiRKAT-O(HMP) | 5.10% | 4.58% | 5.08% | 4.46% | 4.92% | 4.66% |
| | multiMiRKAT-O(ACAT) | 6.08%* | 5.76% | 6.14%* | 5.52% | 6.02%* | 5.92% |
| Optimal test | **multiMiAT(MinP+HMP)** | **4.68%** | **5.34%** | **4.72%** | **4.80%** | **4.96%** | **4.88%** |
| | multiMiAT(MinP+ACAT) | 5.54% | 6.38%* | 5.66% | 5.30% | 5.70% | 5.40% |
| | multiMiAT(HMP) | 4.68% | 5.44% | 4.88% | 4.58% | 5.04% | 4.90% |
| | multiMiAT(ACAT) | 6.32%* | 7.26%* | 6.4%* | 5.94% | 6.84%* | 6.48%* |

Table S5: Empirical type I error rates of microbiome regression-based kernel individual tests and local omnibus tests using the baseline category logit model.

| M | Balance | Kernel | Individual tests and local omnibus tests | | | | | | | |
|---|---|---|---|---|---|---|---|---|---|---|
| | | | $\mathbf{K}_{\mathrm{BC}}$ | $\mathbf{K}_{\mathrm{u}}$ | $\mathbf{K}_{0}$ | $\mathbf{K}_{0.25}$ | $\mathbf{K}_{0.5}$ | $\mathbf{K}_{0.75}$ | $\mathbf{K}_{\mathrm{w}}$ | $\mathbf{K}_{\mathrm{omni}}$ |
| Sample size $n = 100$ | | | | | | | | | | |
| Ordinal | ✓ | LK | 4.90% | 4.82% | 5.08% | 5.04% | 5.22% | 4.98% | 5.04% | 5.24% |
| | | GK | 4.84% | 4.88% | 5.08% | 5.08% | 5.24% | 4.96% | 5.02% | 5.38% |
| | | LaK | 4.78% | 4.76% | 5.14% | 5.04% | 5.10% | 4.82% | 4.86% | 5.20% |
| | × | LK | 4.66% | 5.00% | 5.04% | 5.12% | 4.84% | 4.86% | 4.62% | 5.12% |
| | | GK | 4.66% | 5.02% | 5.06% | 5.08% | 4.84% | 4.86% | 4.60% | 5.06% |
| | | LaK | 4.74% | 5.08% | 4.98% | 4.90% | 4.90% | 4.80% | 4.70% | 5.04% |
| Nominal | | LK | 5.12% | 4.62% | 5.14% | 4.94% | 5.04% | 5.28% | 5.06% | 5.32% |
| | | GK | 5.16% | 4.68% | 5.18% | 4.94% | 5.00% | 5.26% | 5.02% | 5.42% |
| | | LaK | 5.16% | 4.78% | 5.10% | 4.96% | 5.02% | 5.24% | 5.16% | 5.16% |
| Sample size $n = 200$ | | | | | | | | | | |
| Ordinal | ✓ | LK | 4.40% | 4.80% | 4.68% | 4.58% | 4.60% | 4.26% | 4.54% | 4.68% |
| | | GK | 4.38% | 4.74% | 4.84% | 4.58% | 4.66% | 4.18% | 4.56% | 4.62% |
| | | LaK | 4.38% | 4.76% | 4.86% | 4.70% | 4.46% | 4.46% | 4.36% | 4.64% |
| | × | LK | 5.04% | 5.30% | 5.16% | 5.10% | 5.00% | 5.18% | 5.20% | 5.42% |
| | | GK | 5.08% | 5.42% | 5.14% | 5.16% | 5.06% | 5.22% | 5.22% | 5.52% |
| | | LaK | 4.96% | 5.18% | 5.10% | 5.08% | 4.94% | 4.98% | 4.86% | 5.30% |
| Nominal | | LK | 4.94% | 5.06% | 5.44% | 5.26% | 5.94% | 5.56% | 5.40% | 5.28% |
| | | GK | 5.00% | 5.04% | 5.34% | 5.22% | 5.86% | 5.54% | 5.46% | 5.38% |
| | | LaK | 4.92% | 5.16% | 5.50% | 5.18% | 5.80% | 5.60% | 5.22% | 5.26% |

Table S6: Empirical type I error rates of microbiome regression-based kernel individual tests and local omnibus tests using the cumulative link model.

| Category | Balance | Kernel | Individual tests and local omnibus tests | | | | | | | |
|---|---|---|---|---|---|---|---|---|---|---|
| | | | $\mathbf{K}_{\mathrm{BC}}$ | $\mathbf{K}_{\mathrm{u}}$ | $\mathbf{K}_{0}$ | $\mathbf{K}_{0.25}$ | $\mathbf{K}_{0.5}$ | $\mathbf{K}_{0.75}$ | $\mathbf{K}_{\mathrm{w}}$ | $\mathbf{K}_{\mathrm{omni}}$ |
| Sample size $n = 100$ | | | | | | | | | | |
| Ordinal | ✓ | LK | 4.92% | 4.70% | 4.90% | 5.08% | 5.12% | 4.94% | 4.86% | 5.12% |
| | | GK | 5.04% | 4.86% | 4.96% | 5.06% | 5.10% | 4.80% | 4.84% | 5.12% |
| | | LaK | 5.08% | 4.86% | 4.98% | 5.02% | 5.02% | 4.78% | 4.82% | 5.26% |
| | × | LK | 4.80% | 4.92% | 5.10% | 4.98% | 4.94% | 4.82% | 4.74% | 4.94% |
| | | GK | 4.78% | 4.98% | 5.02% | 4.96% | 5.00% | 4.84% | 4.78% | 4.86% |
| | | LaK | 4.98% | 5.20% | 5.16% | 5.02% | 4.92% | 4.90% | 4.68% | 5.00% |
| Nominal | | LK | 5.48% | 5.16% | 5.14% | 5.16% | 5.12% | 5.16% | 5.04% | 5.14% |
| | | GK | 5.50% | 5.18% | 5.04% | 5.20% | 5.14% | 5.06% | 5.02% | 5.10% |
| | | LaK | 5.42% | 5.04% | 4.96% | 5.16% | 4.92% | 5.10% | 5.18% | 5.02% |
| Sample size $n = 200$ | | | | | | | | | | |
| Ordinal | ✓ | LK | 4.40% | 4.64% | 4.92% | 4.50% | 4.54% | 4.26% | 4.32% | 4.88% |
| | | GK | 4.26% | 4.74% | 4.80% | 4.52% | 4.52% | 4.30% | 4.46% | 4.76% |
| | | LaK | 4.30% | 4.86% | 4.84% | 4.60% | 4.52% | 4.36% | 4.18% | 4.60% |
| | × | LK | 4.96% | 5.28% | 5.30% | 5.12% | 4.88% | 5.36% | 5.30% | 5.18% |
| | | GK | 5.04% | 5.38% | 5.32% | 5.28% | 5.00% | 5.36% | 5.34% | 5.08% |
| | | LaK | 4.86% | 5.14% | 5.22% | 5.24% | 4.78% | 5.20% | 4.96% | 5.10% |
| Nominal | | LK | 4.74% | 4.88% | 5.16% | 5.26% | 5.74% | 5.26% | 5.44% | 4.72% |
| | | GK | 4.58% | 5.08% | 5.22% | 5.12% | 5.60% | 5.20% | 5.50% | 4.84% |
| | | LaK | 4.78% | 5.16% | 5.26% | 5.08% | 5.58% | 5.56% | 5.44% | 4.80% |

#### 2.2 Analysis for $p$ values of our methods in real data

Table S7: $P$ values of microbiome regression-based kernel individual tests and local omnibus tests in diverse datasets. $*$ represents a significant association (i.e., $p$ value is below the significance level of 5%).

| Model | Kernel | Individual tests and local omnibus tests | | | | | | | |
|---|---|---|---|---|---|---|---|---|---|
| | | $\mathbf{K}_{\text{BC}}$ | $\mathbf{K}_{\text{u}}$ | $\mathbf{K}_{0}$ | $\mathbf{K}_{0.25}$ | $\mathbf{K}_{0.5}$ | $\mathbf{K}_{0.75}$ | $\mathbf{K}_{\text{w}}$ | $\mathbf{K}_{\text{omni}}$ |
| Dataset A: The association between gut microbiome and colorectal cancer | | | | | | | | | |
| bcl | LK | 0.076 | **0.006*** | 0.058 | 0.121 | 0.235 | 0.300 | 0.264 | **0.023*** |
| | GK | 0.087 | **0.007*** | 0.060 | 0.129 | 0.254 | 0.328 | 0.299 | **0.027*** |
| | LaK | 0.122 | **0.008*** | 0.072 | 0.158 | 0.384 | 0.539 | 0.591 | **0.027*** |
| clm | LK | 0.065 | **0.001*** | **0.038*** | 0.096 | 0.214 | 0.307 | 0.314 | **0.005*** |
| | GK | 0.085 | **0.005*** | 0.051 | 0.113 | 0.266 | 0.436 | 0.515 | **0.017*** |
| | LaK | 0.128 | **0.007*** | 0.052 | 0.135 | 0.368 | 0.575 | 0.707 | **0.024*** |
| Dataset B: The association between microbiome and *Clostridium difficile* infections | | | | | | | | | |
| bcl | LK | **<0.001*** | **<0.001*** | **<0.001*** | **<0.001*** | **<0.001*** | **<0.001*** | **<0.001*** | **<0.001*** |
| | GK | **<0.001*** | **<0.001*** | **<0.001*** | **<0.001*** | **<0.001*** | **<0.001*** | **<0.001*** | **<0.001*** |
| | LaK | **<0.001*** | **<0.001*** | **<0.001*** | **<0.001*** | **<0.001*** | **<0.001*** | **<0.001*** | **<0.001*** |
| clm | LK | **<0.001*** | **<0.001*** | **<0.001*** | **<0.001*** | **<0.001*** | **<0.001*** | **<0.001*** | **<0.001*** |
| | GK | **<0.001*** | **<0.001*** | **<0.001*** | **<0.001*** | **<0.001*** | **<0.001*** | **<0.001*** | **<0.001*** |
| | LaK | **<0.001*** | **<0.001*** | **<0.001*** | **<0.001*** | **<0.001*** | **<0.001*** | **<0.001*** | **<0.001*** |

Table S8: $P$ values of score tests, microbiome regression-based kernel global omnibus tests (i.e., multiMiRKAT-N and multiMiRKAT-O), and optimal tests (i.e., multiMiAT) in diverse datasets. $*$ represents a significant association (i.e., $p$ value is below the significance level of 5%).

| Category | Method | Dataset A | Dataset B |
|---|---|---|---|
| Individual tests | Score test | 0.084 | **<0.001*** |
| Omnibus tests | MiRKAT-MCN | **0.040*** | **<0.001*** |
| | MiRKAT-MCO | **0.023*** | **<0.001*** |
| | multiMiRKAT-N | **0.029*** | **<0.001*** |
| | multiMiRKAT-O | **0.006*** | **<0.001*** |
| Optimal tests | multiMiAT | **0.020*** | **<0.001*** |

#### 2.3 Analysis for $p$ values of microbiome-based association tests in real data

Table S9: $P$ values of the microbiome-based association tests for any pair of statuses from the three statuses. $*$ represents a significant association (i.e., $p$ value is below the significance level of 5%). Group 1, 2 and 3 represent health/nondiarrhea, adenoma/diarrhea and carcinoma/case, respectively.

| Method | Dataset A | | | Dataset B | | |
|---|---|---|---|---|---|---|
| | Group 1 and 2 | Group 1 and 3 | Group 2 and 3 | Group 1 and 2 | Group 1 and 3 | Group 2 and 3 |
| aMiSPU | 0.062 | 0.104 | **0.039*** | **0.001*** | **0.001*** | **0.006*** |
| MiHC | **0.007*** | 0.139 | 0.564 | **<0.001*** | **<0.001*** | **0.043*** |
| OMiAT | **0.020*** | **0.030*** | 0.141 | **<0.001*** | **<0.001*** | **<0.001*** |
| OMiRKAT | 0.060 | **0.019*** | 0.082 | **0.001*** | **0.001*** | **0.003*** |

Table S10: $P$ values of the pairwise, adjacent pairwise and baseline pairwise analysis of microbiome-based microbiome tests. $*$ represents a significant association (i.e., $p$ value is below the significance level of 5%).

| Category | Methods | Dataset A | Dataset B |
|---|---|---|---|
| aMiSPU | Pairwise analysis | 0.117 | **0.003*** |
| | Adjacent pairwise analysis | 0.078 | **0.002*** |
| | Baseline pairwise analysis | 0.078 | **0.002*** |
| MiHC | Pairwise analysis | **0.020*** | **<0.001*** |
| | Adjacent pairwise analysis | **0.013*** | **<0.001*** |
| | Baseline pairwise analysis | 0.278 | **<0.001*** |
| OMiAT | Pairwise analysis | 0.060 | **<0.001*** |
| | Adjacent pairwise analysis | **0.040*** | **<0.001*** |
| | Baseline pairwise analysis | 0.060 | **<0.001*** |
| OMiRKAT | Pairwise analysis | 0.056 | **0.003*** |
| | Adjacent pairwise analysis | 0.119 | **0.002*** |
| | Baseline pairwise analysis | **0.037*** | **0.002*** |

### 3 Supplementary Figures

#### 3.1 Analysis for the hyperparameter $\sigma$ and choice of kernel functions

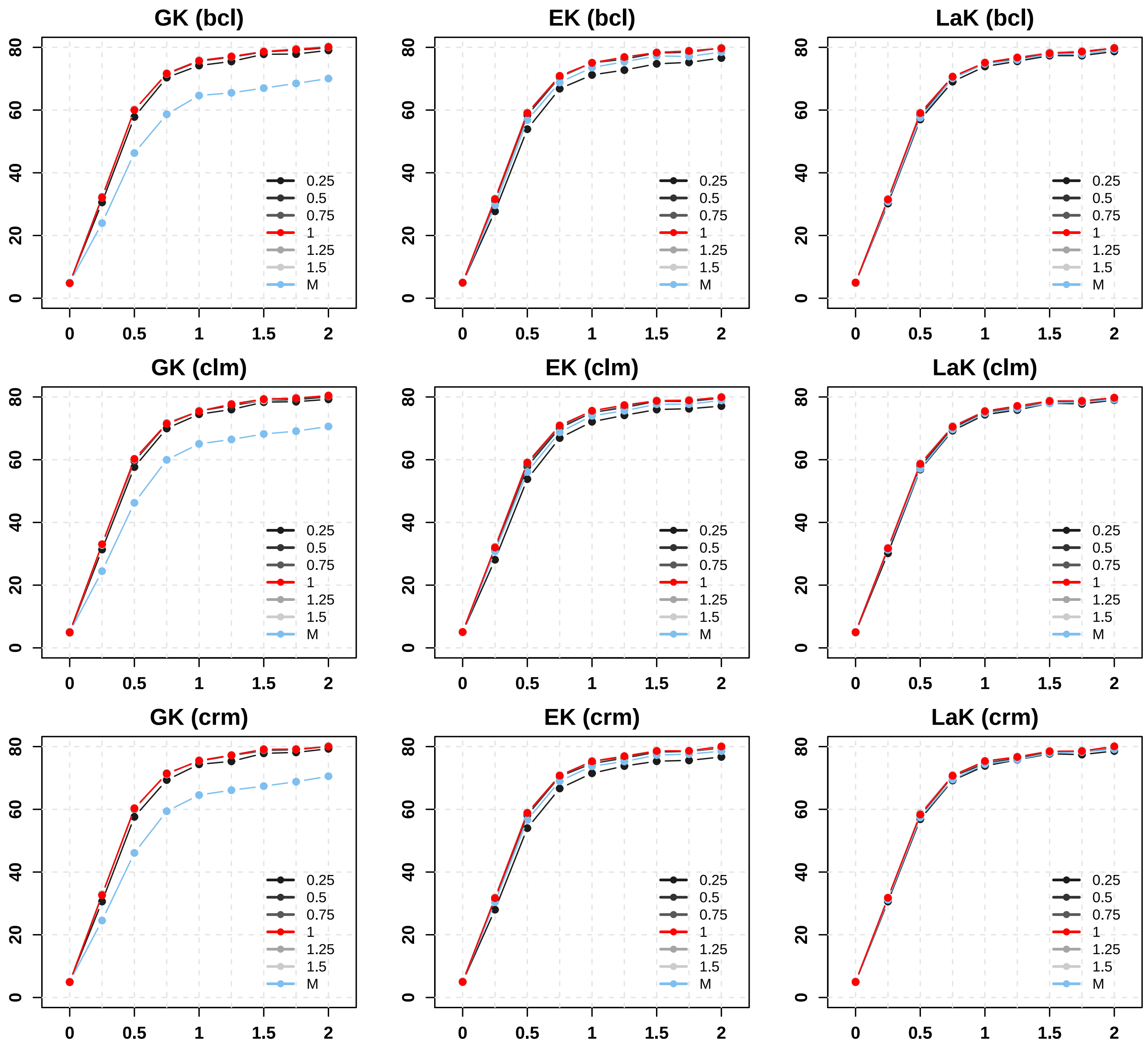

Figure S1: Comparative analysis of the powers for microbiome regression-based kernel local omnibus test with diverse $\sigma$ in scenario 1a ($n = 100$). (A) Fitting baseline category logit model. (B) Fitting cumulative link model. (C) Fitting cumulative link model. Here, the local omnibus tests integrate the individual tests with eight distances (i.e., Jaccard dissimilarity [13], Bray-Curtis dissimilarity, UniFrac distance, generalized UniFrac with $\alpha = 0$, 0.25, 0.5, 0.75, and weighted UniFrac.) Here, "M" denotes the mean of distance matrix $\mathbf{D}$ [10].

#### 3.2 Empirical type I error rates and power analysis of our individual tests

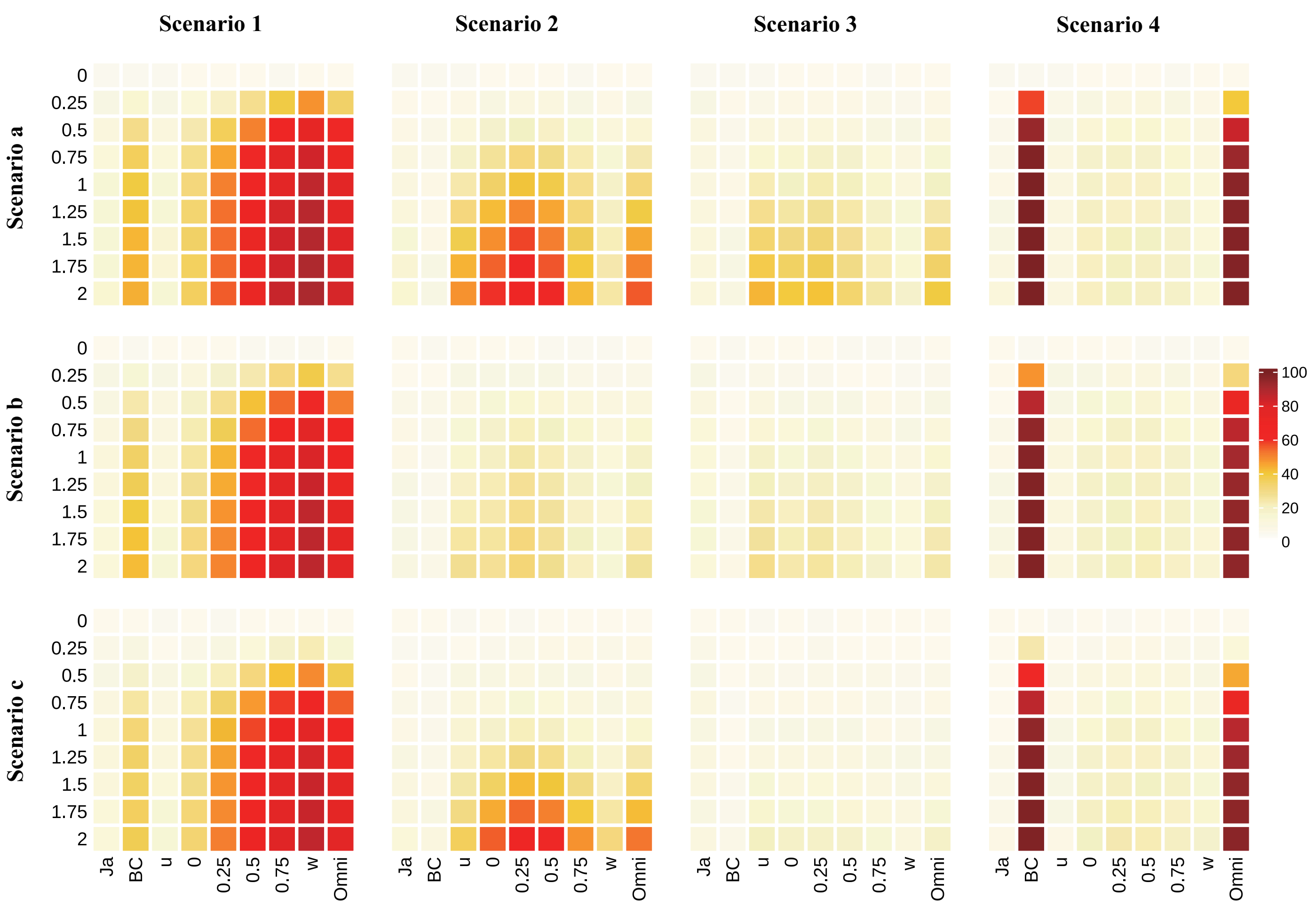

Figure S2: Comparison of the type I error rates and powers among microbiome regression-based kernel individual tests and local omnibus test with diverse distances under diverse scenarios ($n = 100$). Here, we adopt the linear kernel and the baseline category logit model.

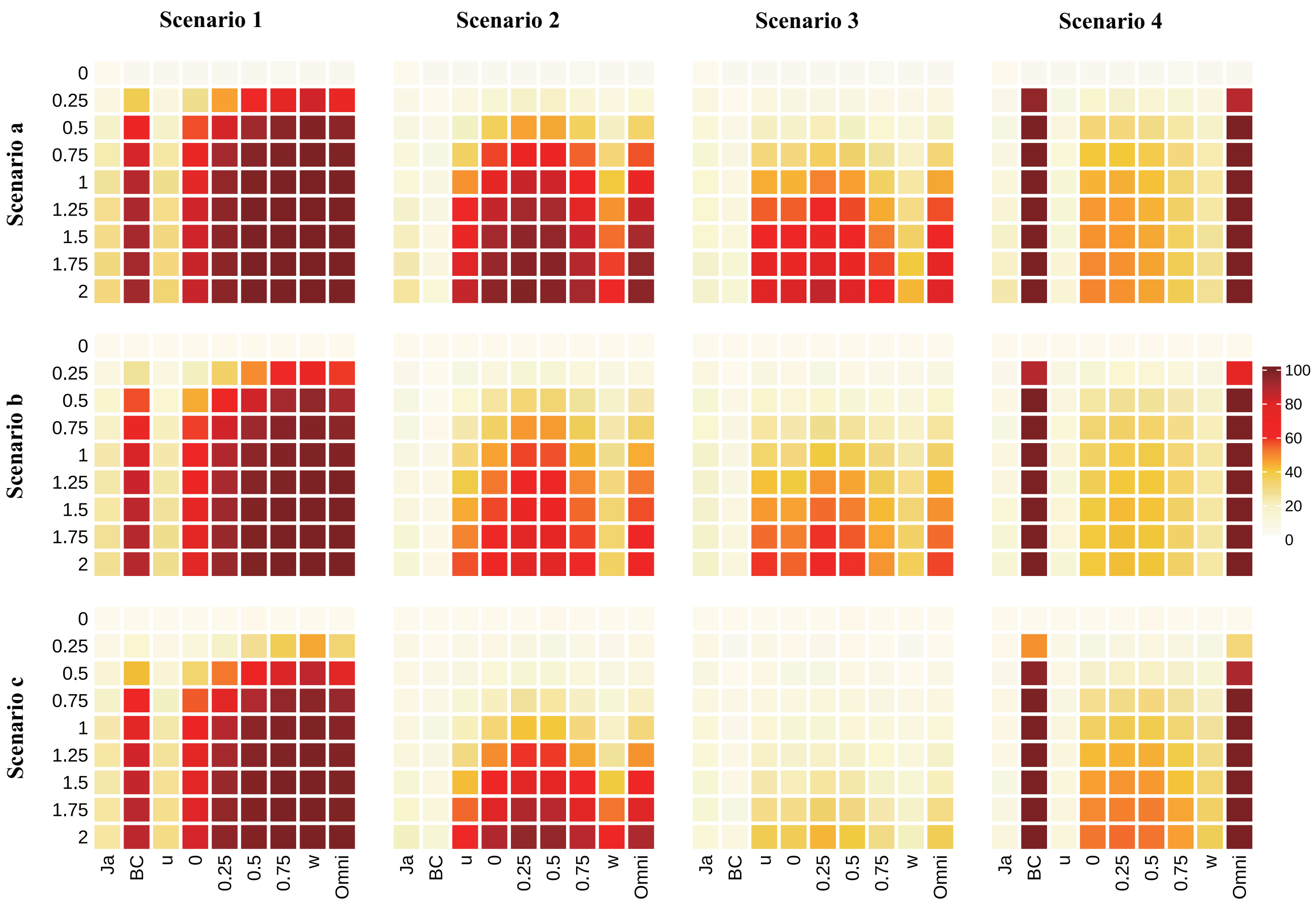

Figure S3: Comparison of the type I error rates and powers among microbiome regression-based kernel individual tests and local omnibus test with diverse distances under diverse scenarios ($n = 200$). Here, we adopt the Gaussian kernel and the baseline category logit model.

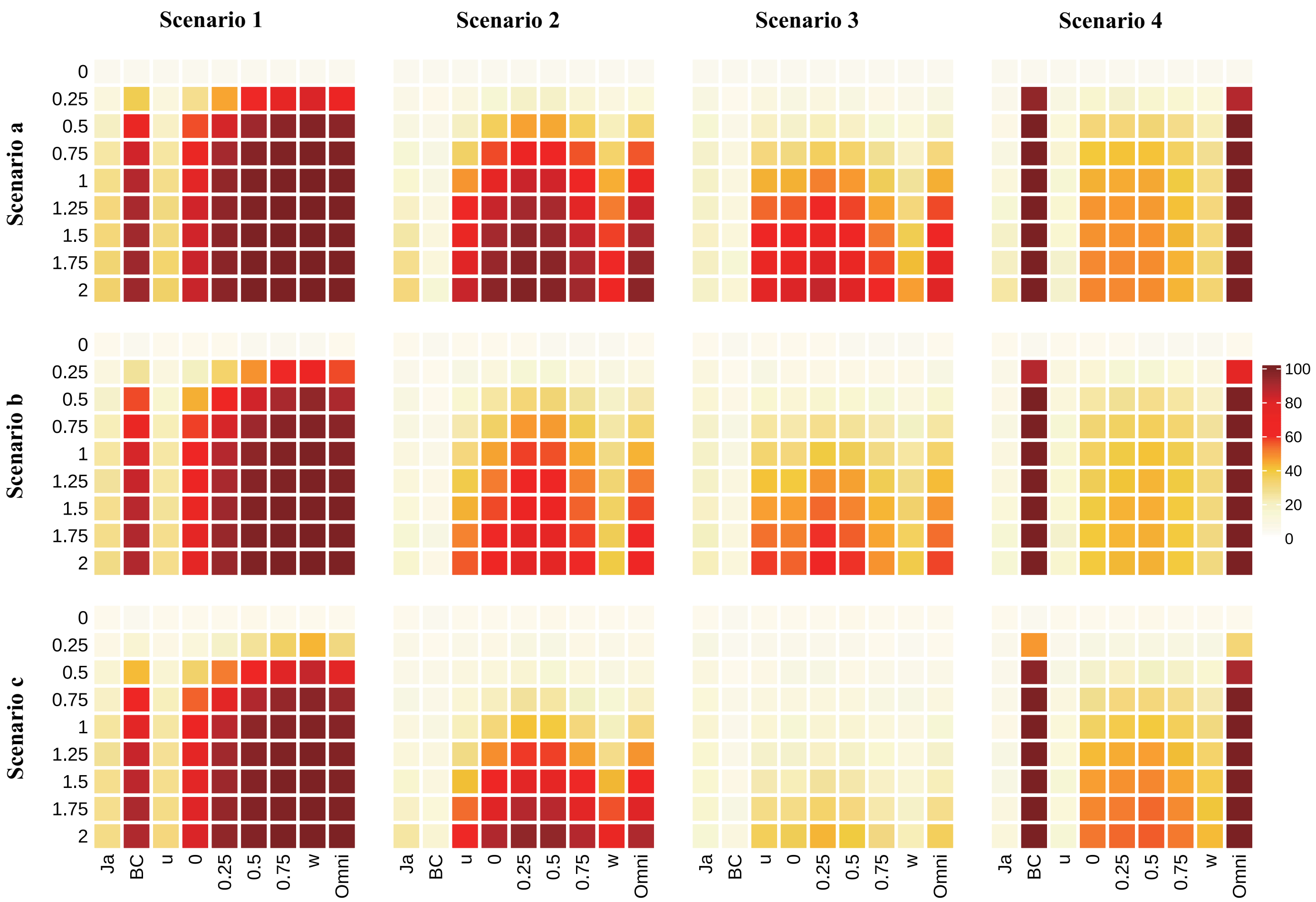

Figure S4: Comparison of the type I error rates and powers among microbiome regression-based kernel individual tests and local omnibus test with diverse distances under diverse scenarios ($n = 200$). Here, we adopt the Laplacian kernel and the baseline category logit model.

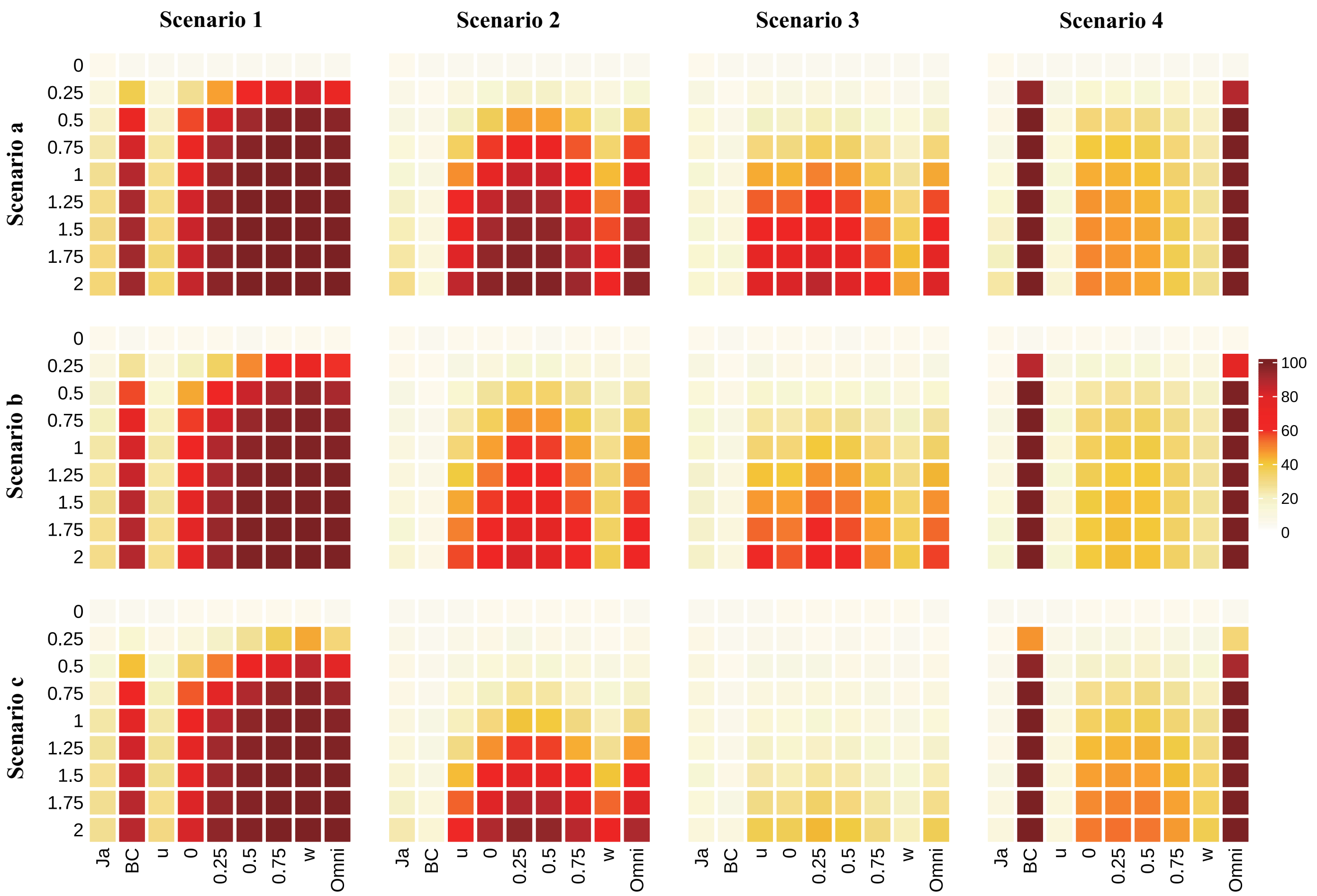

Figure S5: Comparison of the type I error rates and powers among microbiome regression-based kernel individual tests and local omnibus test with diverse distances under diverse scenarios ($n = 200$). Here, we adopt the linear kernel and the cumulative link model.

##### 3.3 Analysis for the choice of ordinal multinomial logit models

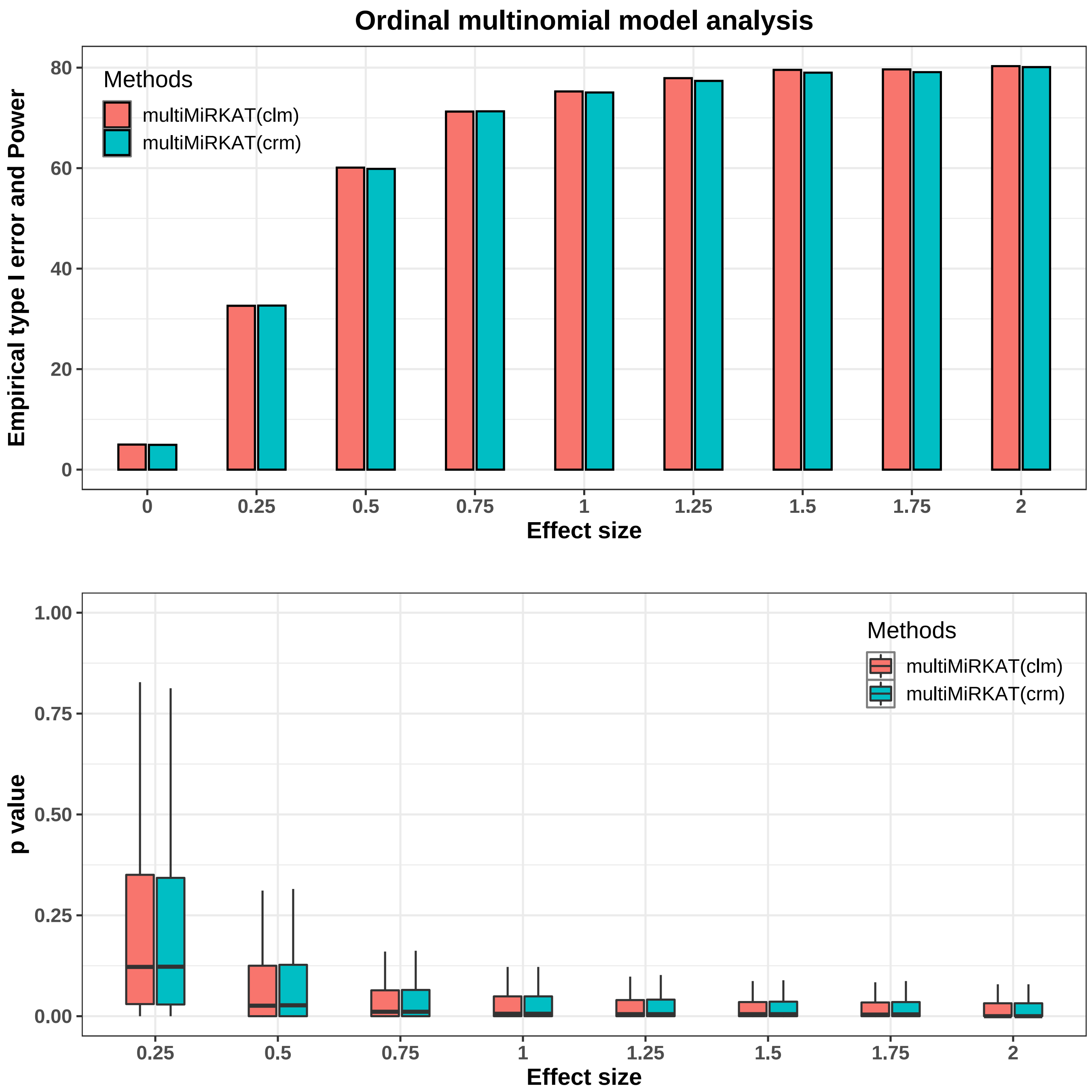

Figure S6: Comparative analysis of microbiome regression-based kernel omnibus tests which are with different ordinal multinomial models (i.e., clm and crm) in scenario 1a ($n = 100$). (A) Comparison of power. (B) Comparison of $p$ value distribution. Here, multiMiRKAT tests also integrate the individual tests with eight distances (i.e., Jaccard dissimilarity, Bray-Curtis dissimilarity, UniFrac distance, generalized UniFrac with $\alpha = 0$, 0.25, 0.5, 0.75, and weighted UniFrac.)

#### 3.4 Analysis for the powers of microbiome regression-based kernel local omnibus tests and global omnibus tests

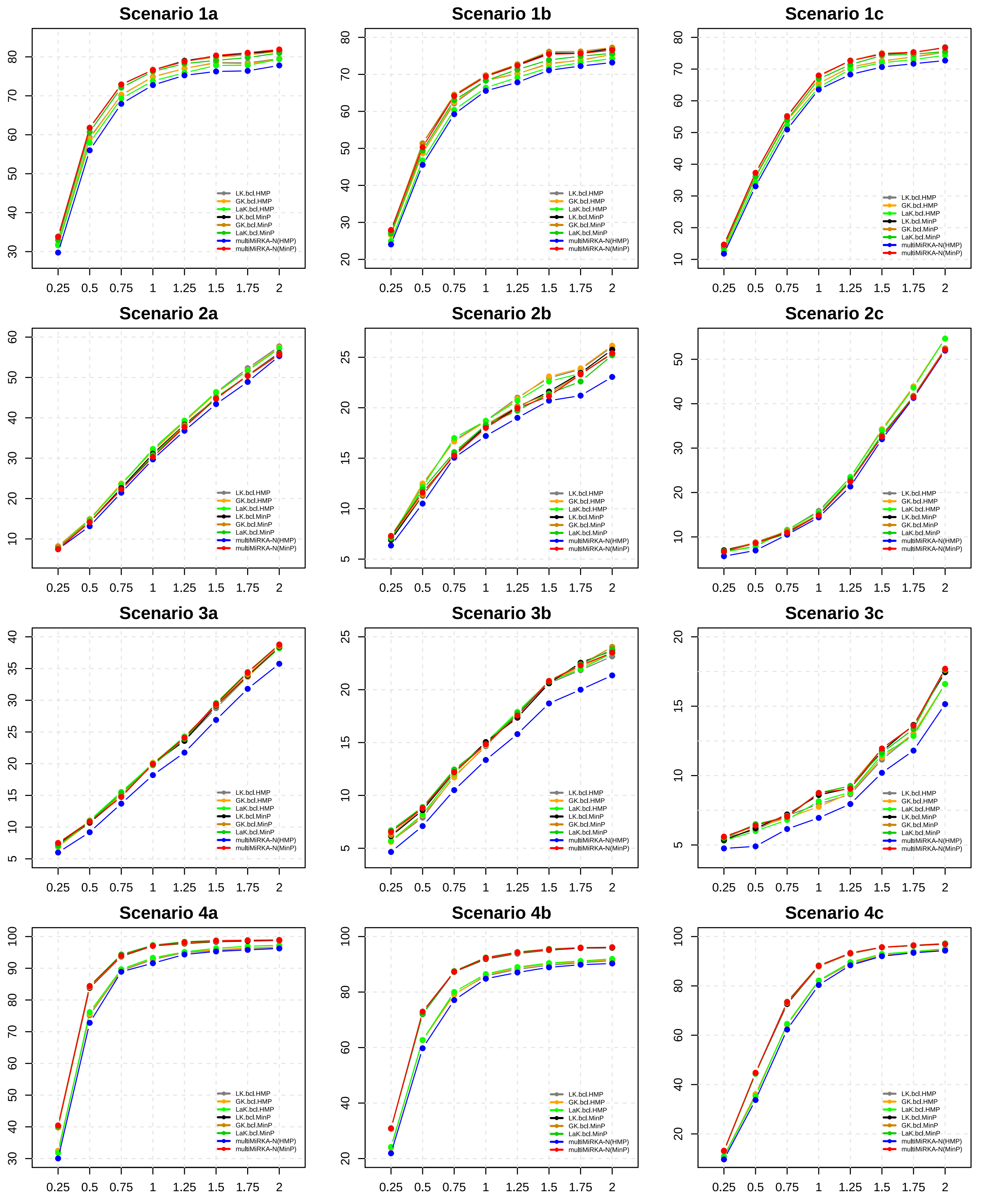

Figure S7: Comparison of the powers among microbiome regression-based kernel local omnibus tests and global omnibus test based on the baseline category logit model (i.e., multiMiRKAT-N) under diverse scenarios ($n = 100$).

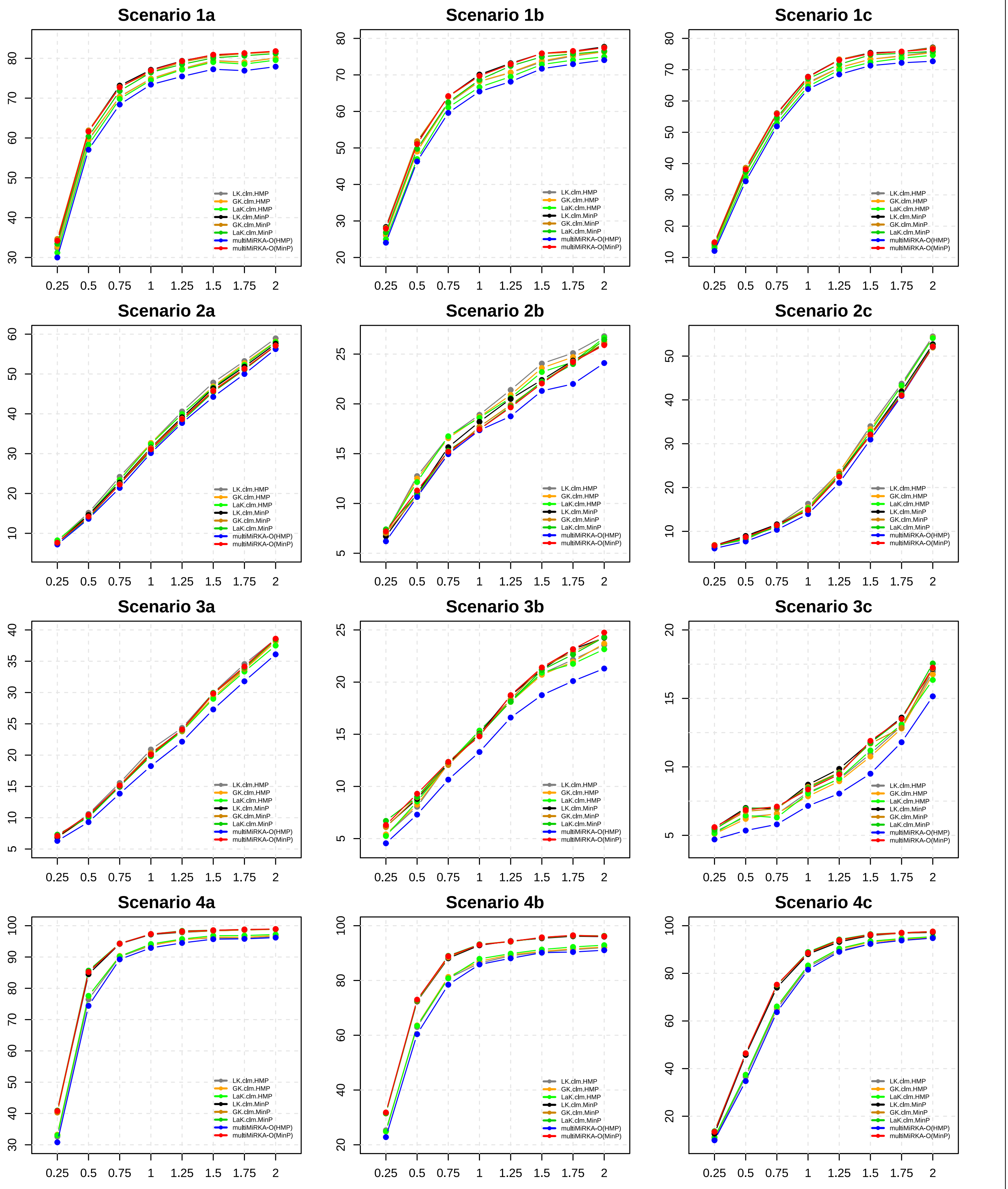

Figure S8: Comparison of the powers among microbiome regression-based kernel local omnibus tests and global omnibus test based on the cumulative link model (i.e., multiMiRKAT-N) in diverse scenarios ($n = 100$).

#### 3.5 Analysis for the powers of combined tests and optimal test

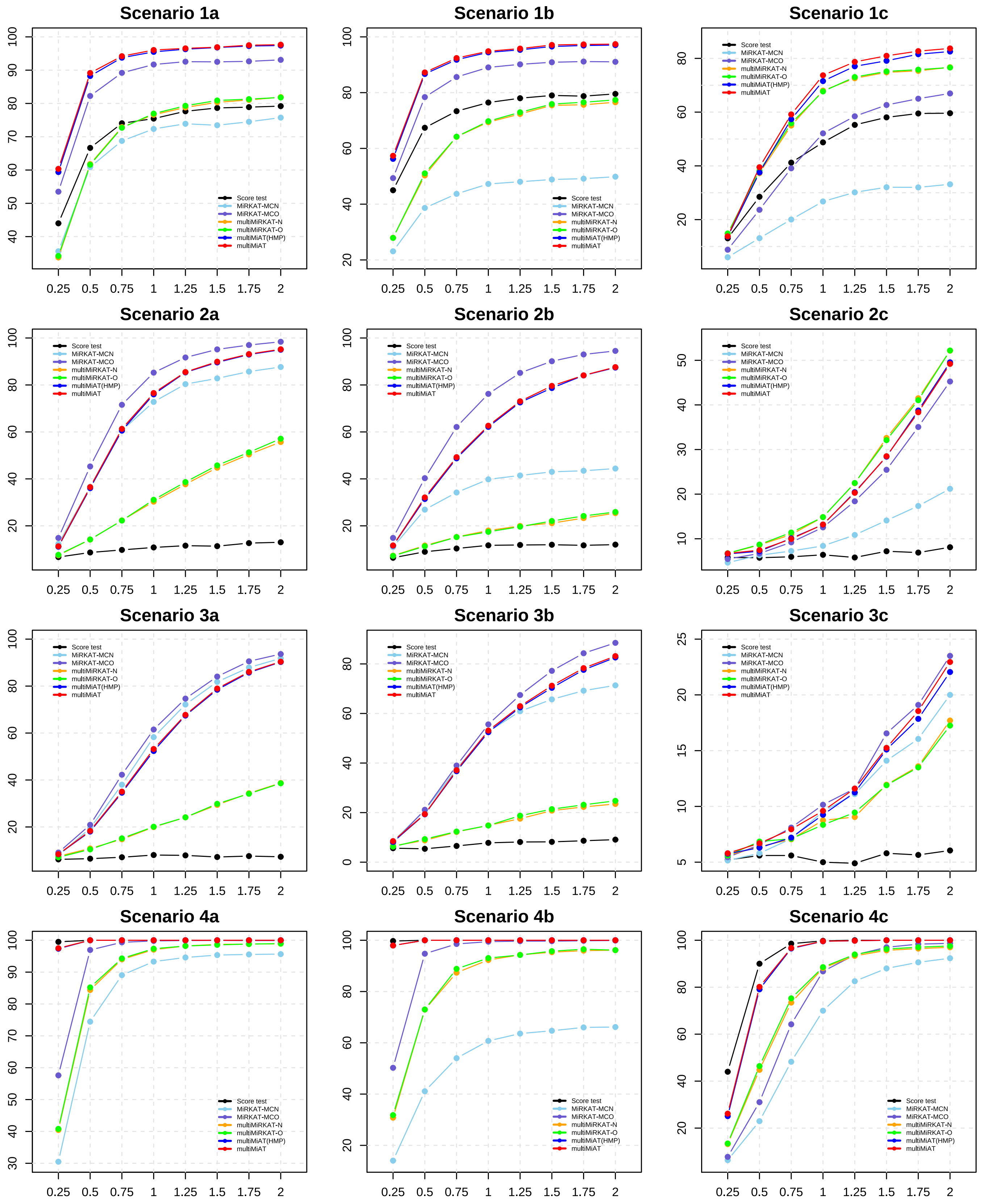

Figure S9: Comparison of the powers among combined tests (i.e., score test, MiRKAT-MCN test, MiRKAT-MCO test, multiMiRKAT-N test, multiMiRKAT-O test) and optimal test (i.e., multiMiAT test) under diverse scenarios ($n = 100$).

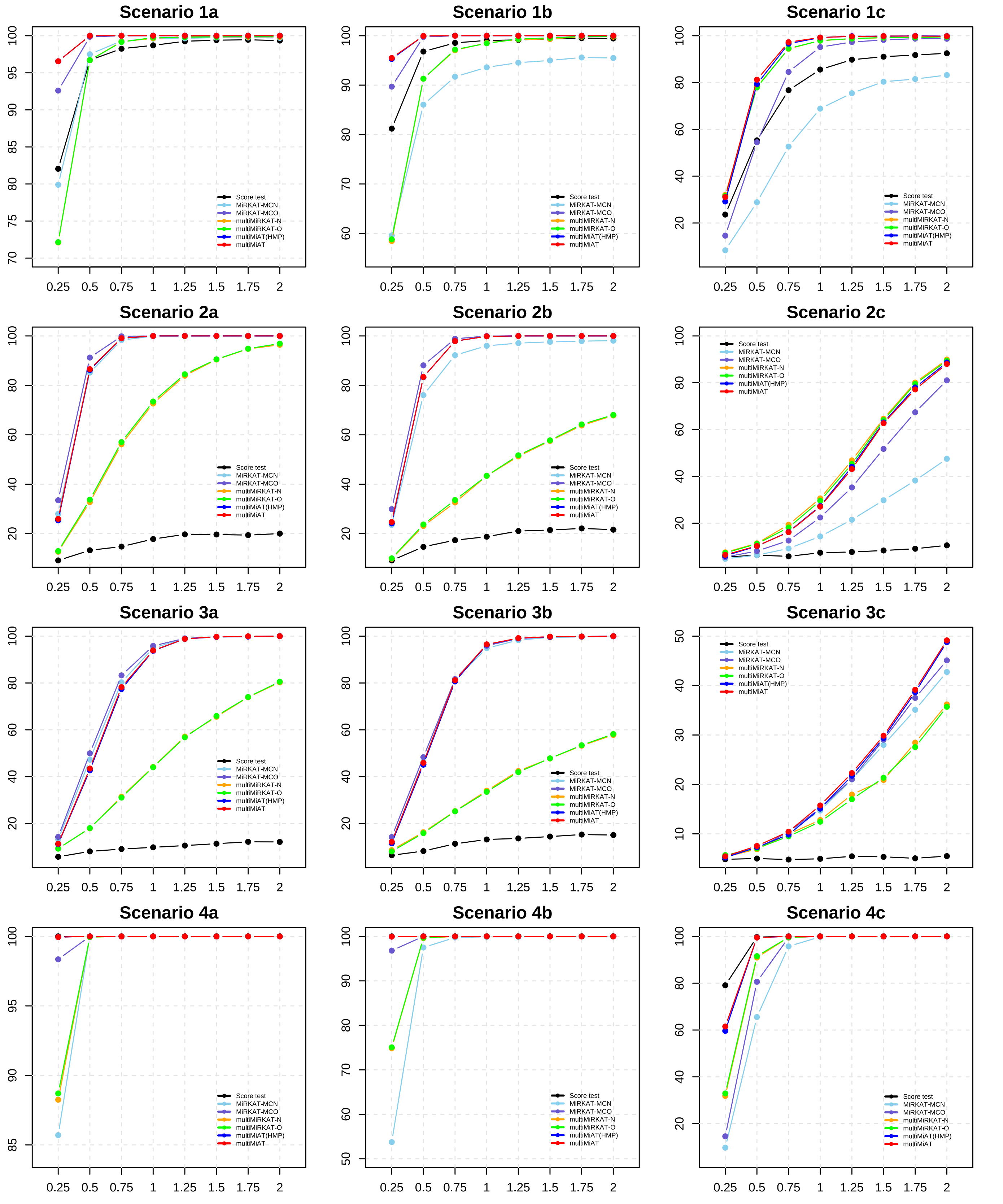

Figure S10: Comparison of the powers among combined tests (i.e., score test, MiRKAT-MCN test, MiRKAT-MCO test, multiMiRKAT-N test, multiMiRKAT-O test) and optimal test (i.e., multiMiAT test) under diverse scenarios ($n = 200$).

##### 3.6 Distribution of minimal $p$ value in pairwise analysis of microbiome-based association tests

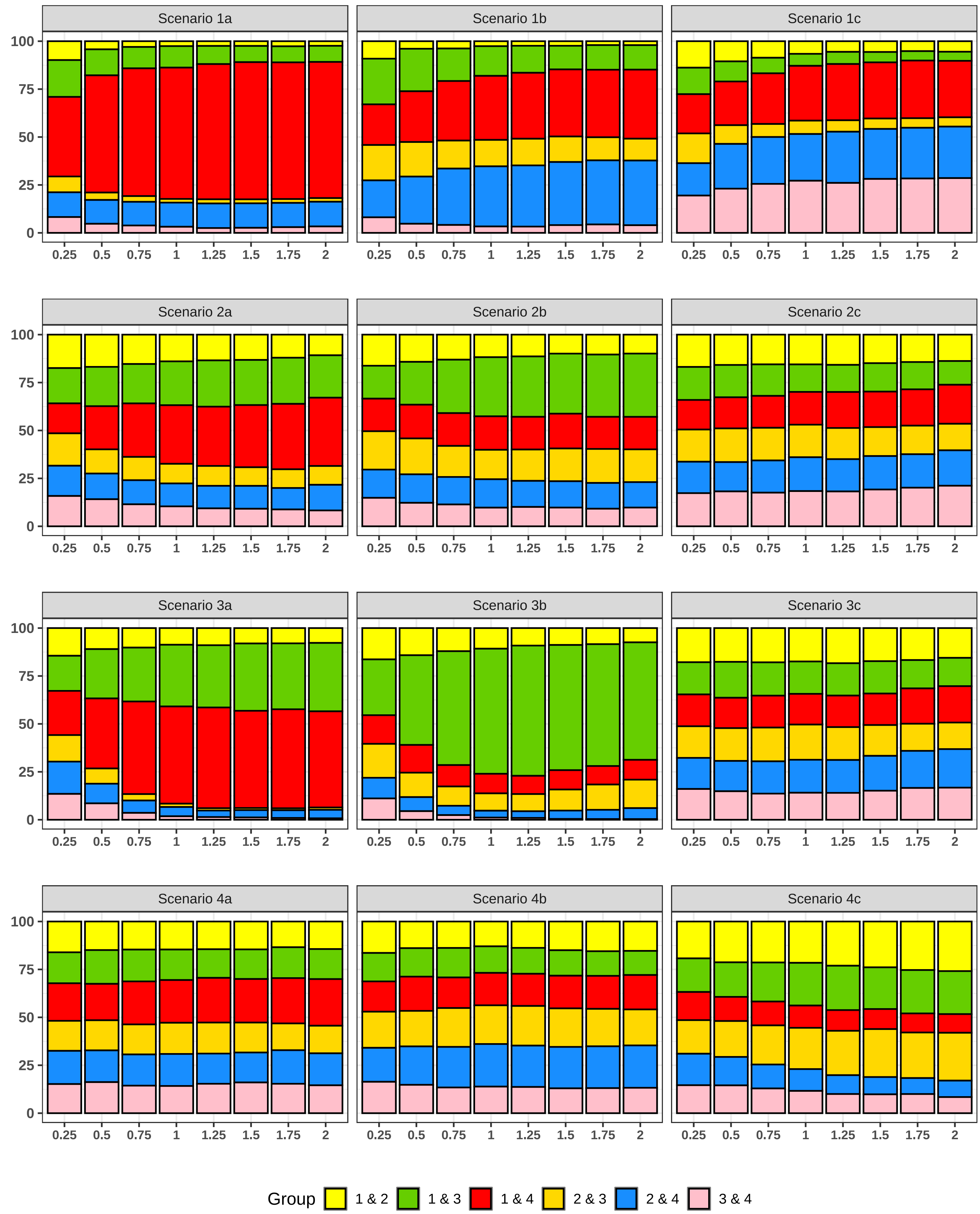

Figure S11: The distribution of minimal $p$ value in $J(J-1)/2$ pairwise tests of aMiSPU ($n = 100$).

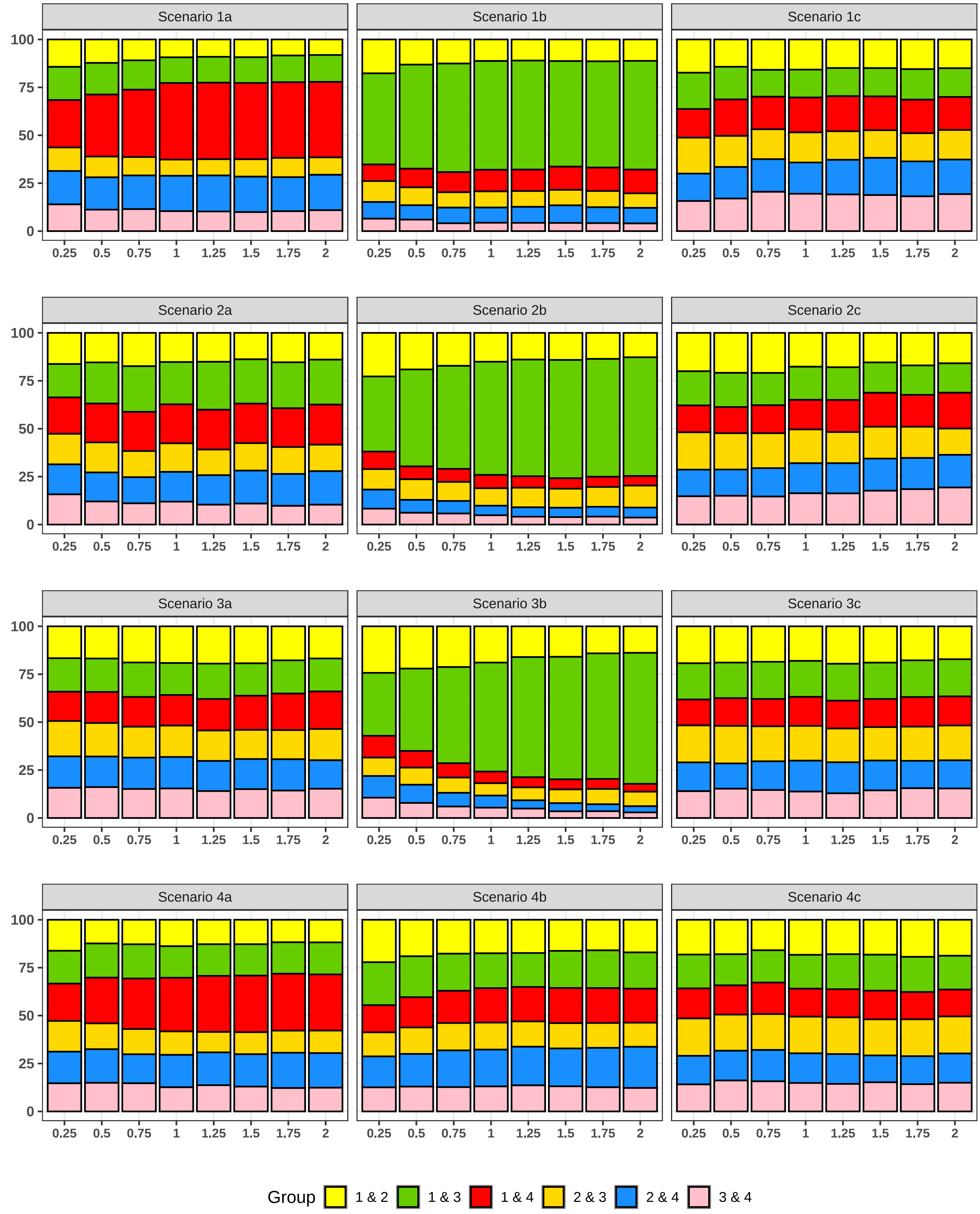

Figure S12: The distribution of minimal $p$ value in $J(J-1)/2$ pairwise tests of MiHC ($n = 100$).

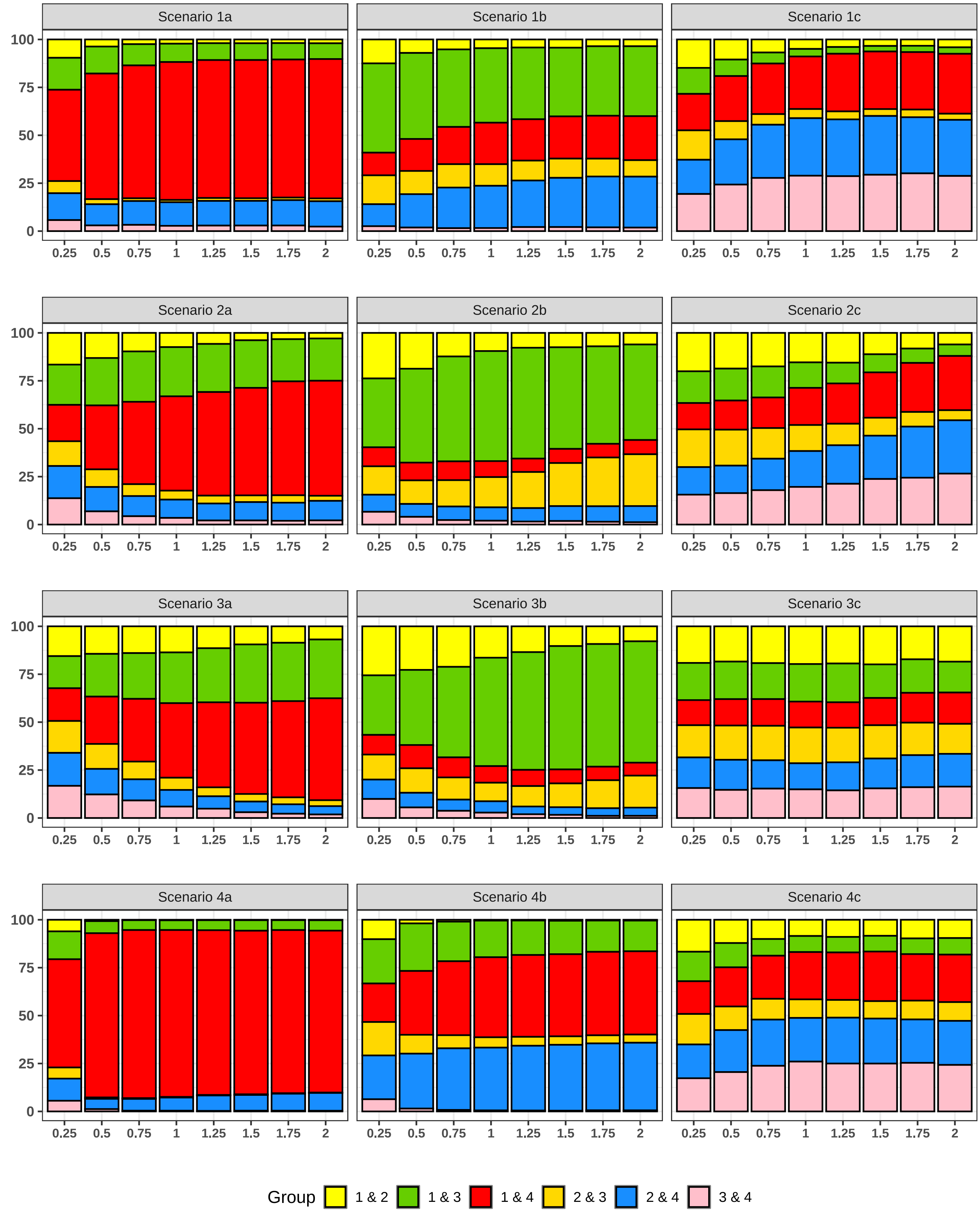

Figure S13: The distribution of minimal $p$ value in $J(J-1)/2$ pairwise tests of OMiAT ($n = 100$).

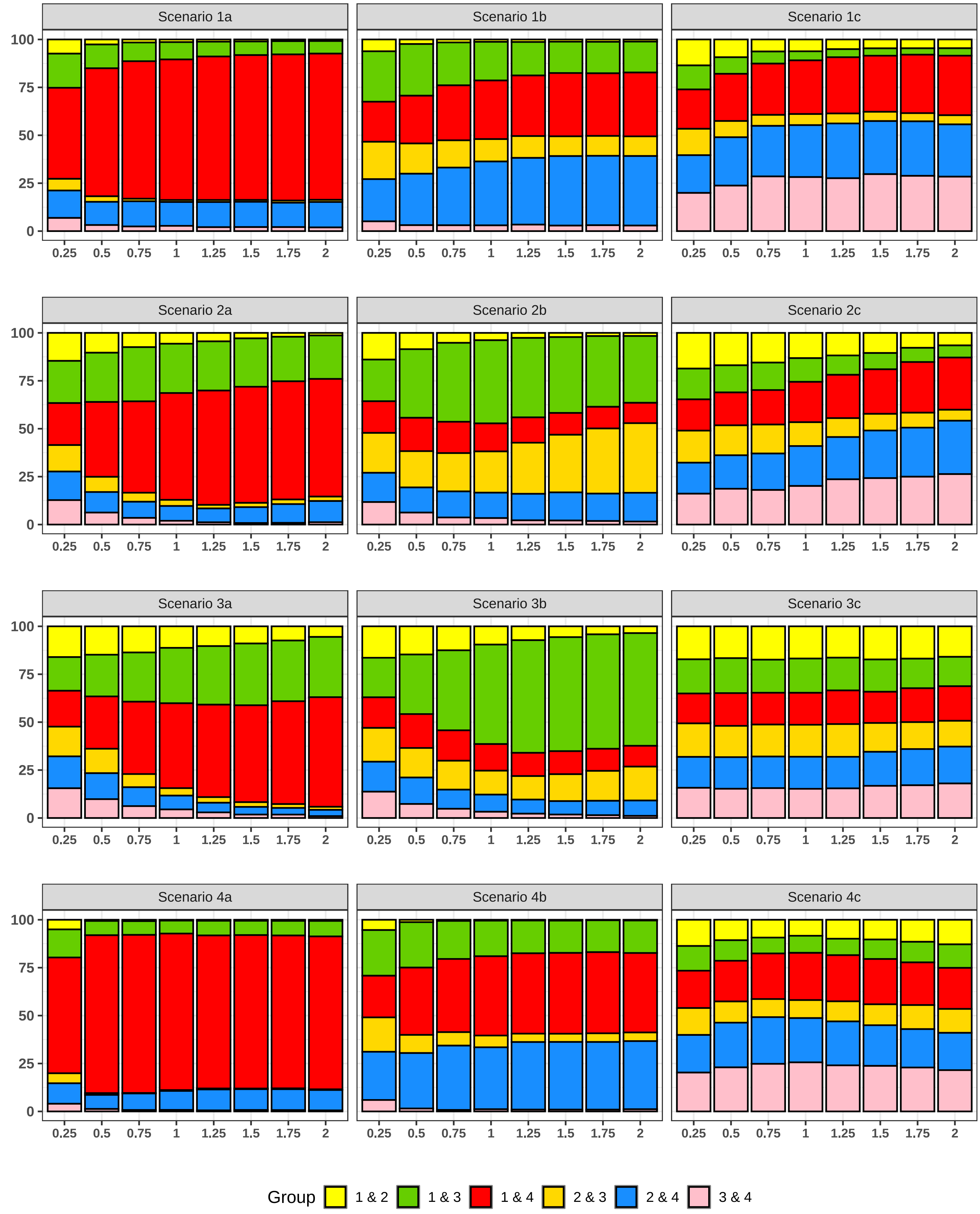

Figure S14: The distribution of minimal $p$ value in $J(J-1)/2$ pairwise tests of OMiRKAT ($n = 100$).

##### 3.7 Analysis for the powers of pairwise, adjacent pairwise and baseline pairwise analysis of microbiome-based microbiome tests

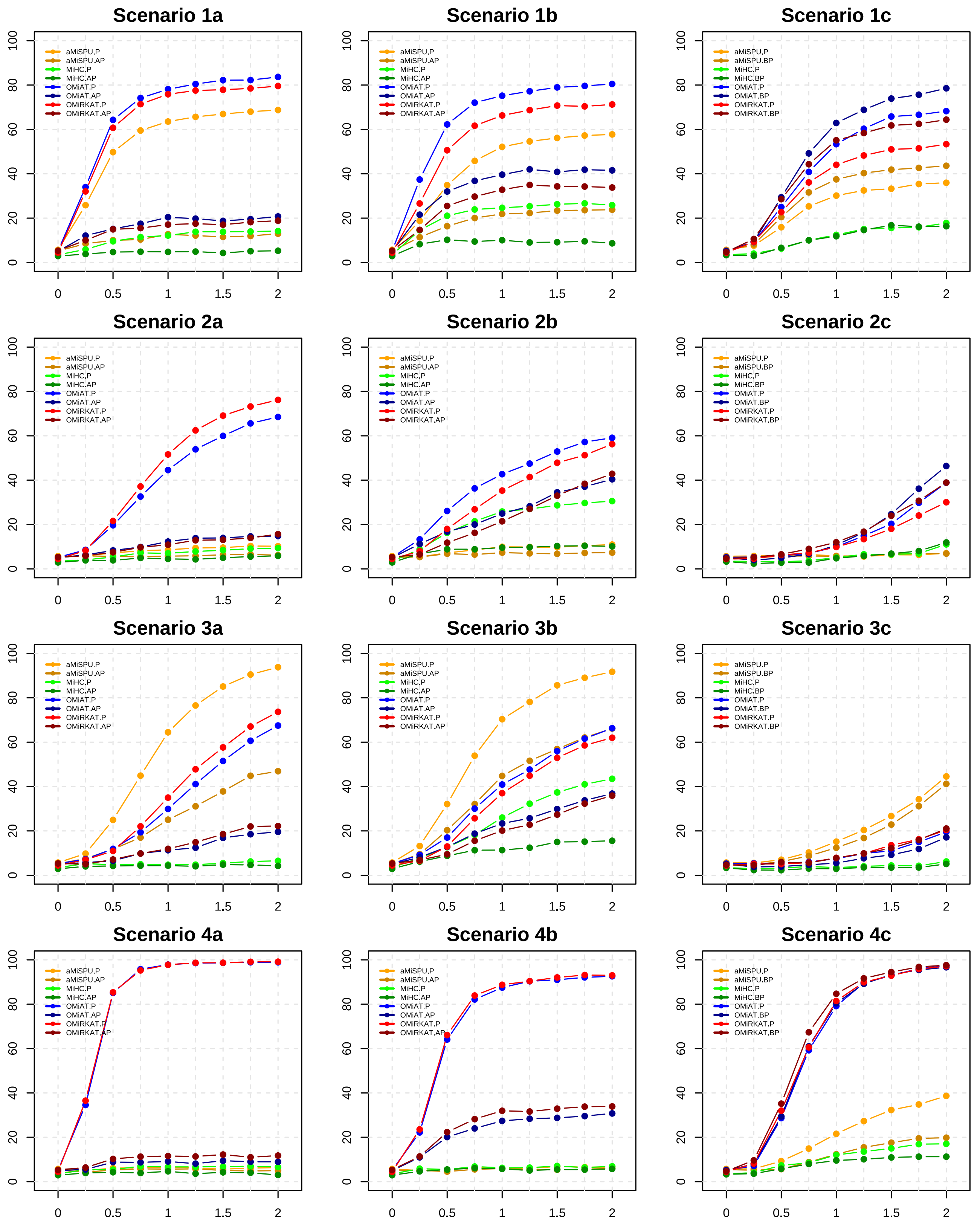

Figure S15: Comparative analysis of the diverse correction strategies of microbiome-based association tests ($n = 100$). Here, "aMiSPU.P", "aMiSPU.AP" and "aMiSPU.BP" represent pairwise, adjacent pairwise and baseline pairwise analysis of aMiSPU, respectively.

#### 3.8 Analysis for the powers of our method and previous methods

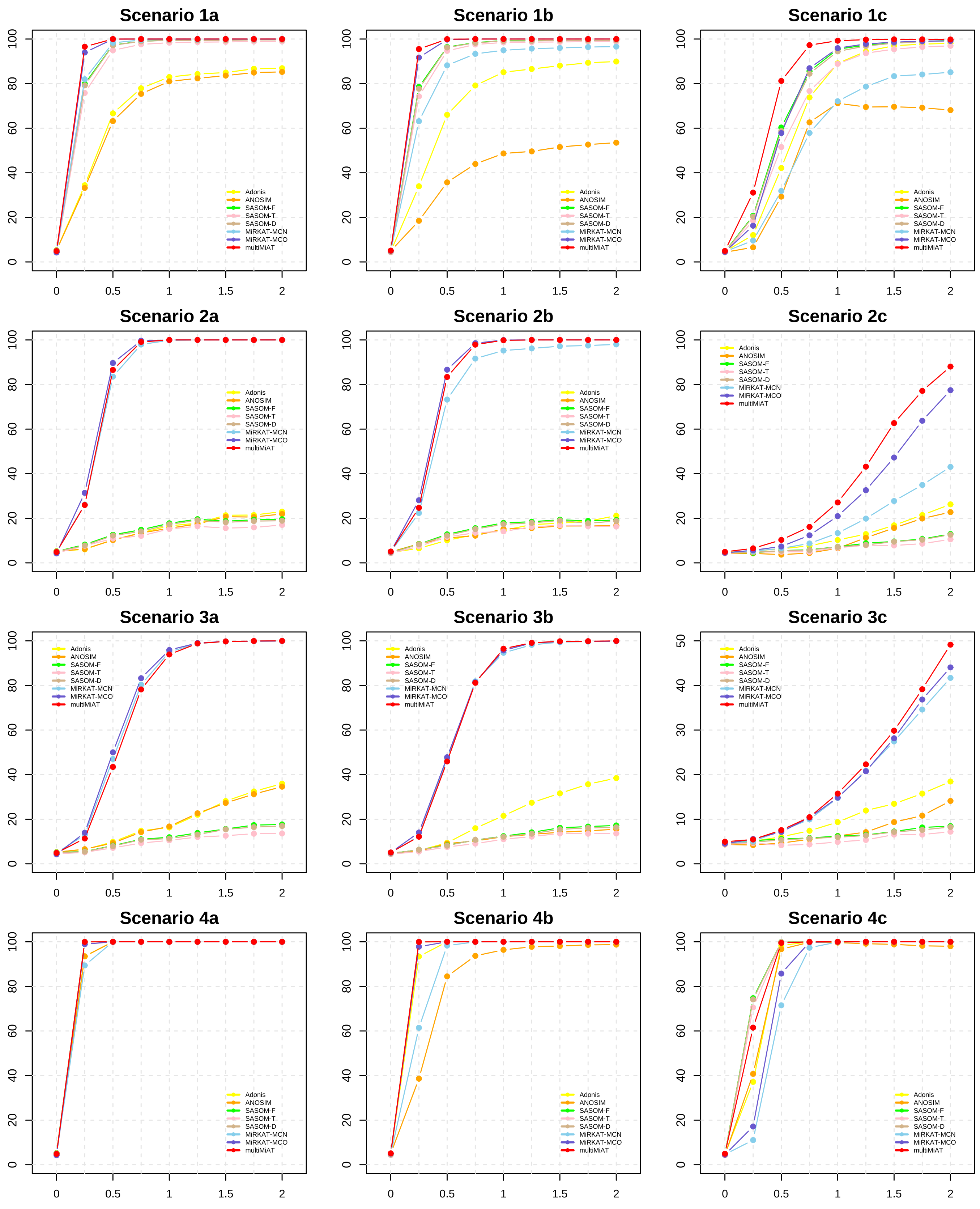

Figure S16: Under diverse scenarios, the comparison of the powers between our method (i.e., multiMiAT) and previous association tests for multicategory outcomes ($n = 200$).

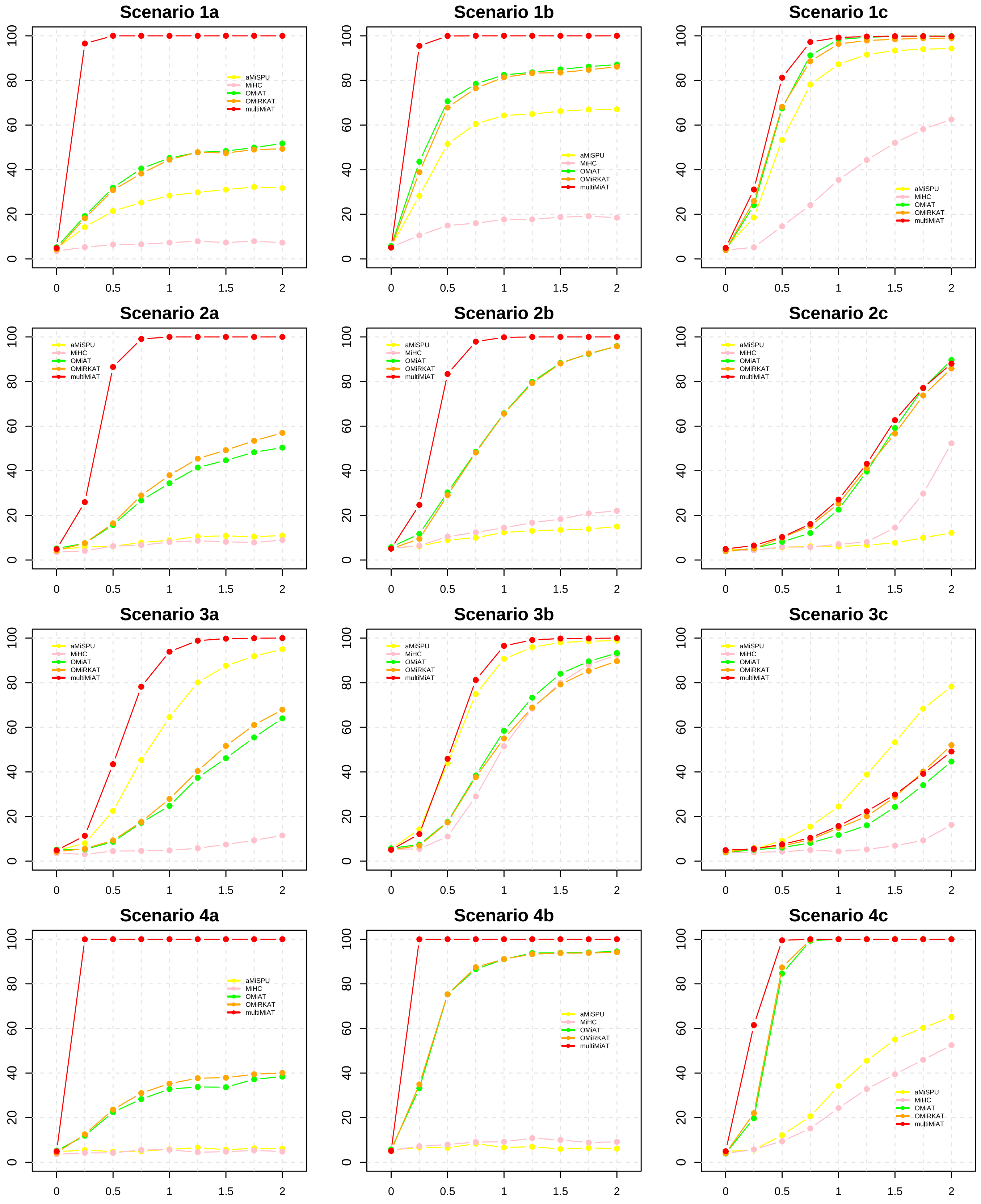

Figure S17: Under diverse scenarios, the comparison of the powers between our method (i.e., multiMiAT) and previous microbiome-based methods ($n = 200$).

##### 3.9 Empirical type I error rate and power analysis for synthesizing all scenarios

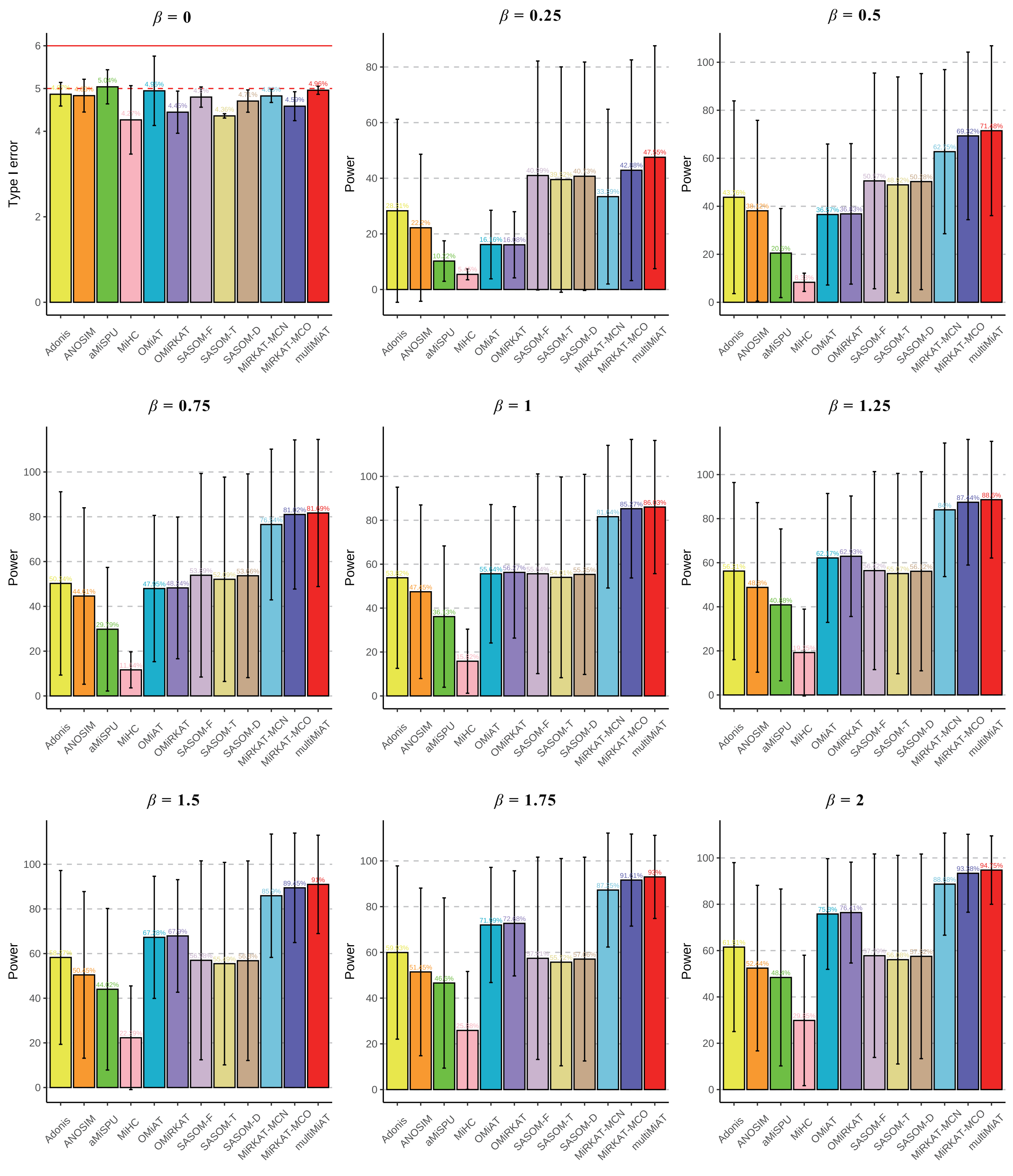

Figure S18: Comparison of empirical type I error rates and powers for synthesizing all scenarios under diverse effect sizes $\beta$ ($n = 200$).

##### 3.10 Principal co-ordinates analysis in diverse development statuses of CRC

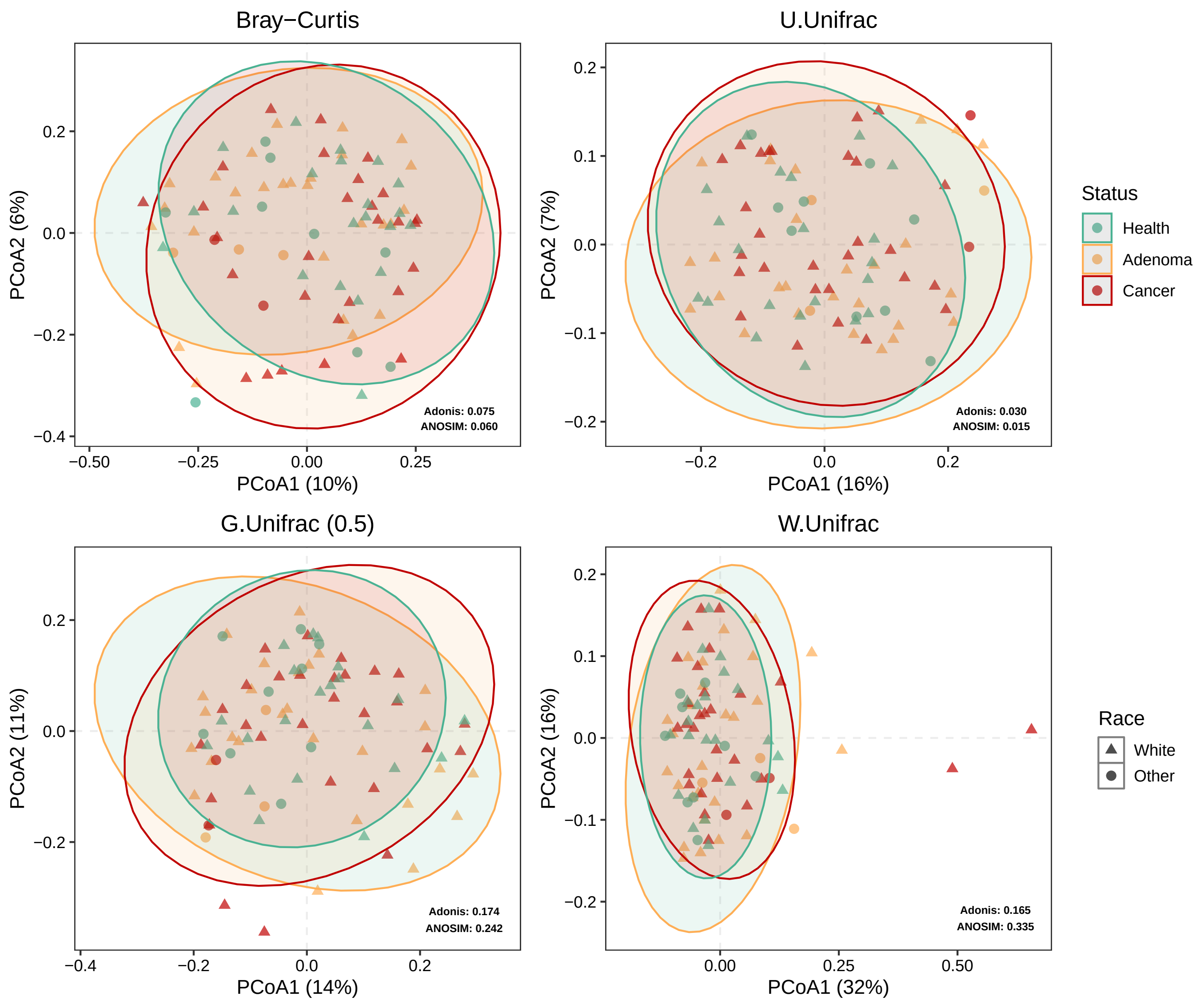

Figure S19: Principal co-ordinates analysis (PCoA) plot based on diverse dissimilarity metrics (i.e., Bray-Curtis dissimilarity, UniFrac distance, GUniFrac distance and WUniFrac distance) between samples in dataset A.

##### 3.11 Difference analysis of gut microbiome in three clinical statuses of CRC development

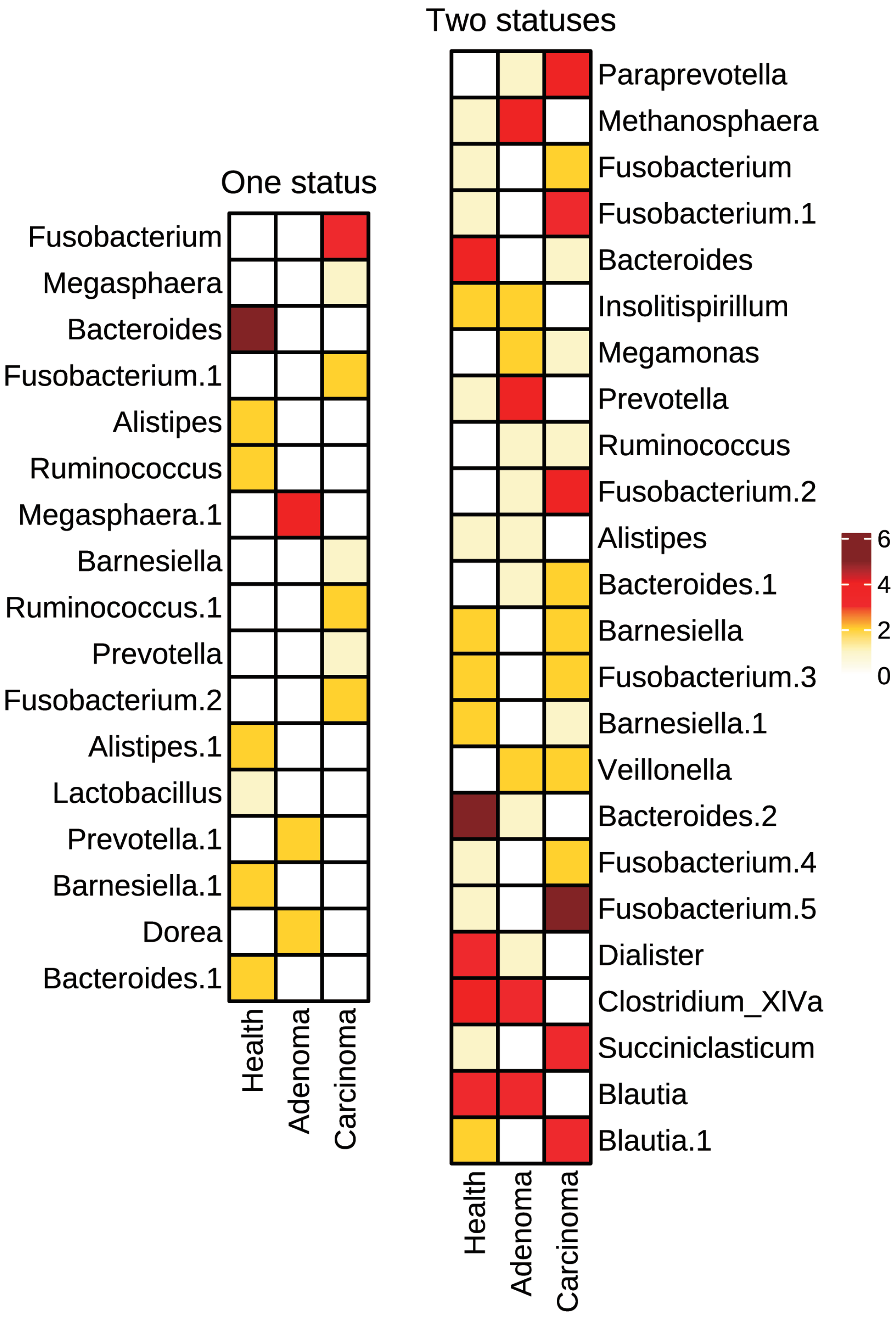

Figure S20: Frequency heatmap of OTU presence in three statuses. OTUs exist in only one status or two statuses.

##### 3.12 Principal co-ordinates analysis in diverse development statuses of CDI

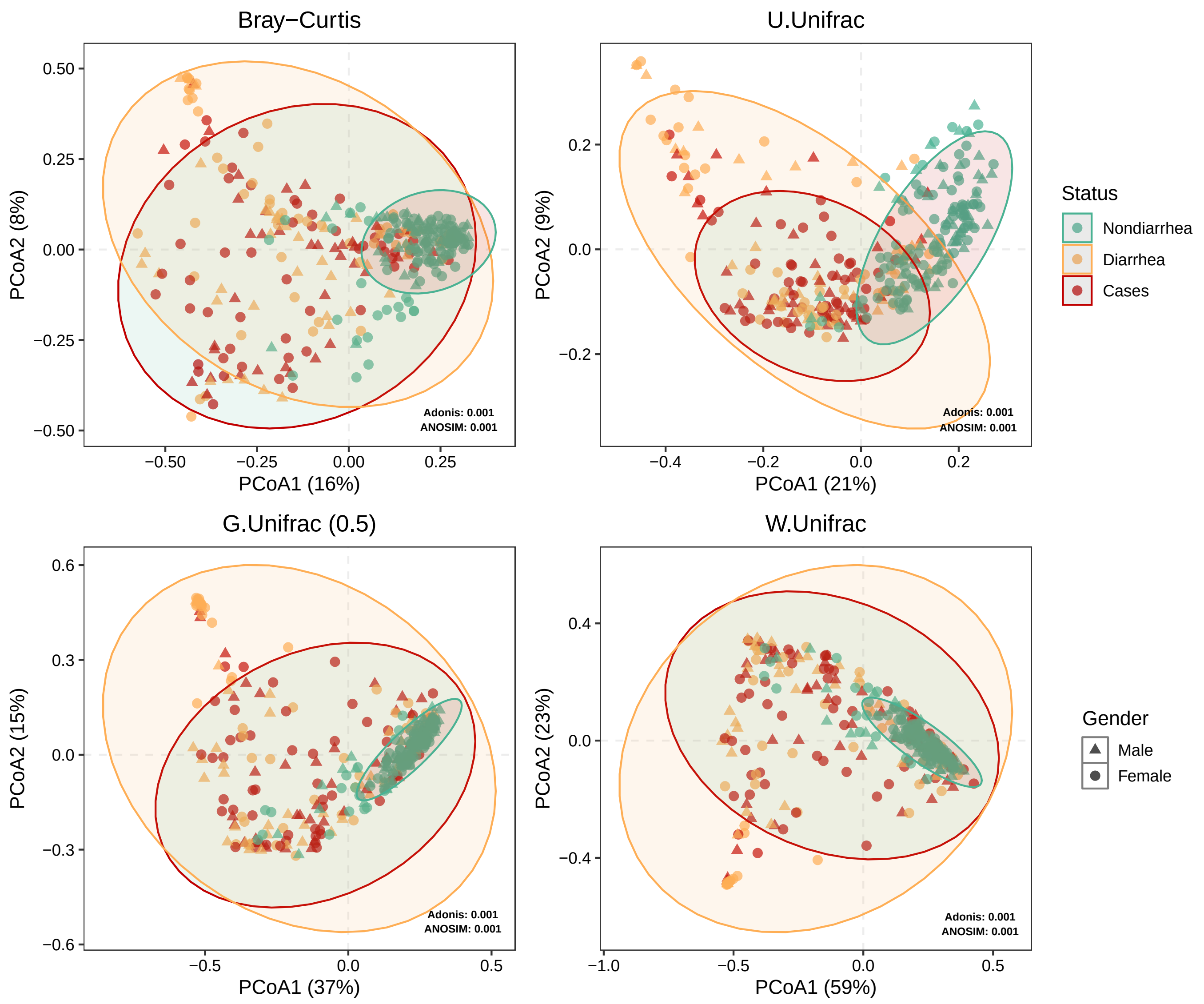

Figure S21: Principal co-ordinates analysis (PCoA) plot based on diverse dissimilarity metrics (i.e., Bray-Curtis dissimilarity, UniFrac distance, GUniFrac distance and WUniFrac distance) between samples in dataset B.

### References

[1] Yee TW, et al. The VGAM Package for Categorical Data Analysis. *Journal of Statistical Software* 2010;**32**(10).

[2] Bray JR, Curtis JT. An Ordination of the Upland Forest Communities of Southern Wisconsin. *Ecological Monographs* 1957;**27**(4):325-349.

[3] Lozupone CA, Knight Rob. UniFrac: a New Phylogenetic Method for Comparing Microbial Communities. *Applied and Environmental Microbiology* 2007;**73**(5):1576-1585.

[4] Lozupone C, et al. Quantitative and qualitative $\beta$ diversity measures lead to different insights into factors that structure microbial communities. *Applied and Environmental Microbiology* 2007;**73**(5):1576-1585.

[5] Chen J, et al. Associating microbiome composition with environmental covariates using generalized UniFrac distances. *Bioinformatics* 2012;**28**(16):2106-2113.

[6] Agarwal D, Zhang NR. Semblance: Anempirical similarity kernel on probability spaces. *Science Advances* 2019;**5**(12):eaau9630.

[7] Zhao N, et al. Testing in microbiome-profiling studies with MiRKAT, the microbiome regression-based kernel association test. *American Journal of Human Genetics* 2015;**96**(5):797-807.

[8] Liu M, et al. A method for subtype analysis with somatic mutations. *Bioinformatics* 2021;**37**(1):50-56.

[9] Jiang Z, et al. MiRKAT-MC: A distance-based microbiome kernel association test with multi-categorical outcomes. *Frontiers in Genetics* 2022;**13**:841764.

[10] Zhan X, et al. A fast small-sample kernel independence test for microbiome community-level association analysis. *Biometrics* 2017;**173**(4):1453-1463.

[11] Wilson DJ. The harmonic mean $p$-value for combining dependent tests. *Proceedings of the National Academy of Sciences* 2019;**116**(4):1195-1200.

[12] Liu Y, et al. ACAT: A fast and powerful p value combination method for rare-variant analysis in sequencing studies. *American Journal of Human Genetics* 2019; **104**(3):410-421.

[13] Jaccard P. The distribution of the flora in the alpine zone. *New Phytologist* 1912;**11**(2):37-50.
